# Skeletal muscle enhancer programming of intrinsic cardiorespiratory fitness

**DOI:** 10.1101/2025.10.02.679855

**Authors:** Anne M. Weitzel, Peter Orchard, Charles R. Evans, Nandini Manickam, Mary K. Treutelaar, Steven L. Britton, Lauren Gerard Koch, Jun Z. Li, Stephen C.J. Parker, Charles F. Burant

**Author notes:** Co-Senior Author (Stephen C.J. Parker, Charles F. Burant).

## Abstract

Cardiorespiratory fitness (CRF) is a heritable trait associated with improved metabolic health and longevity. Integrating 546 transcriptomic and epigenomic profiles from skeletal muscle of 128 genetically heterogeneous rats selectively bred for high and low running capacity (HCR and LCR), a model that mirrors CRF-associated traits in humans, we identify a distributed enhancer architecture underlying their divergent transcriptomes. Line-divergent alleles were concentrated in distal enhancers: those more active in HCR were enriched for ETS-family transcription factor motifs and positively linked to hundreds of mitochondrial oxidative metabolism and angiogenesis genes, whereas those more active in LCR were enriched for Mef2 motifs and negatively linked to these genes. In 426 genotype, gene expression, and chromatin accessibility profiles from an independent HCR×LCR F2 population (n=147), we validated thousands of these genetic effects. Thus, heritable CRF reflects many enhancers cumulatively tuning energy metabolism and oxygen delivery, providing a molecular basis for variation in CRF.

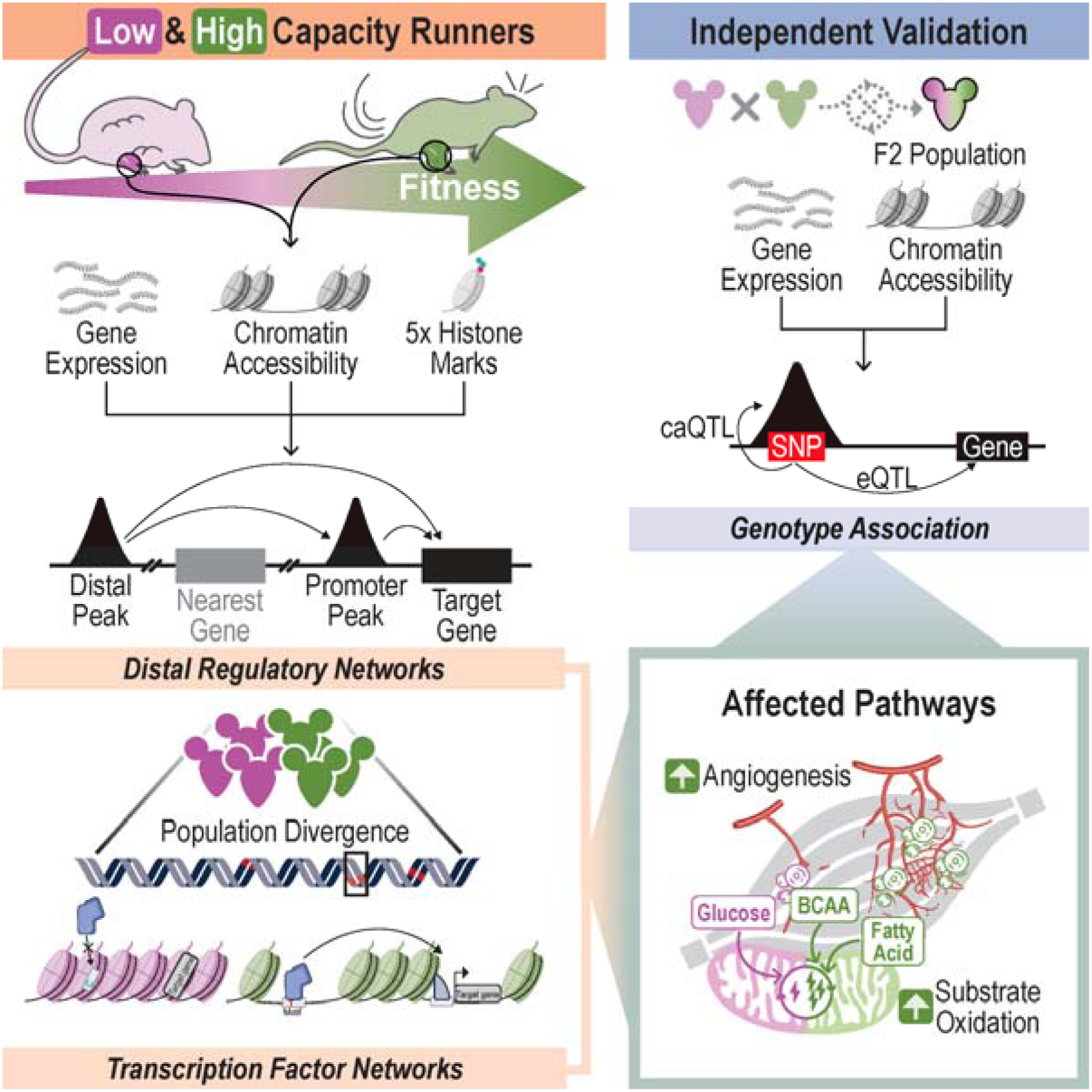

## Introduction

Physical activity is recognized for its positive effect on human health and longevity^1,2^. Conversely, sedentary time independently increases risk of cardiometabolic disease and mortality^1–3^. Independently, cardiorespiratory fitness (CRF), measured by the maximal oxygen uptake (VO_2_max) during a graded exercise test, is a strong predictor of cardiometabolic risk and mortality^4,5^ even in the presence of preexisting cardiovascular disease and its risk factors^6,7^. CRF is a heritable trait with 50-75% of its variation being genetically determined (*intrinsic* CRF)^8–10^. Regular exercise can increase VO_2_max by 8-15% and has shown particularly strong fitness and longevity effects in individuals with low intrinsic CRF^11–13^, though these individuals still face higher risks of cardiometabolic disease and premature death than their non-exercising, high CRF counterparts^12^. While large-scale human population studies have identified genetic variants associated with CRF^14,15^ distinct from those associated with exercise training^16^, the molecular mechanisms underlying the variation in CRF remain poorly defined.

To study intrinsic CRF, we use the High Capacity Runner (HCR) and Low Capacity Runner (LCR) model^17^ derived from the N:NIH heterogeneous rat stock^18^ by selection for VO_2_max, generating two divergent lines^17^. This well-characterized model shows persistent and heritable phenotypes that recapitulate human traits associated with divergent CRF, including sex-independent differences in physiologic measures of cardiometabolic health and longevity that correlate with VO_2_max^19,20^ (Fig. 1a). As seen in humans with high CRF, the HCR line has an increased capacity for fatty acid oxidation and shows upregulation of genes involved in fatty acid and branched chain amino acid metabolism in skeletal muscle^21,22^ and heart^23^. We previously generated an HCR×LCR (F2) cross population whose segregating genotypes produced heritable variation in running capacity and muscle phenotypes^24^ (Fig. 1b). The genetic variants differentiated through selection, along with the F2 population, provide a unique and tractable system to study the skeletal muscle regulatory architecture and its contribution to intrinsic CRF.

**Fig. 1:**
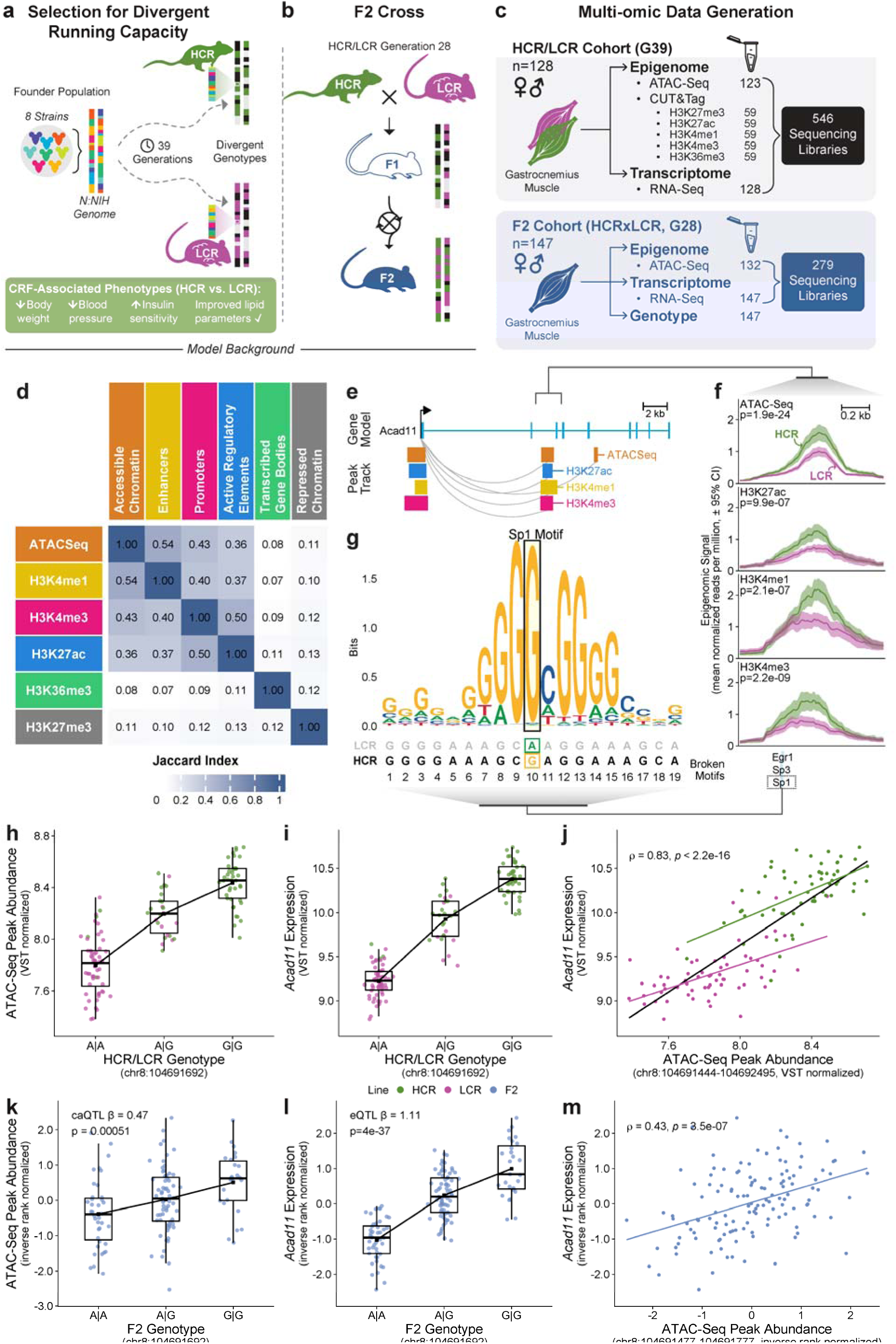
Selection for divergent running capacity presents a regulatory cascade at *Acad11*. (**a**) HCR/LCR model background. Illustration of the genetic mosaic of the N:NIH heterogeneous stock, originating from eight inbred rat strains (distinct colors), that served as the founder population for the HCR and LCR lines. Multiple generations of artificial selection for intrinsic running capacity led to allele fixation and divergence between HCR (green) and LCR (magenta) populations, resulting in distinct genotypic and phenotypic profiles in each line. **(b)** Crossing scheme used for previous generation of the genetically diverse F2 population. First, HCR and LCR lines were crossed to produce an F1 population, illustrated by heterozygous loci inherited from both parental lines. The F1 generation was then randomly intercrossed, while avoiding sibling pairs, to produce the genetically recombinant F2 population containing a mixture of genetic variants derived from both HCR and LCR backgrounds. (**c**) Multi-omic profiling of gastrocnemius muscle from 128 HCR/LCR rats (equal representation of HCR/LCR and sex) and 147 F2 rats (equal representation of sex), including RNA-Seq, ATAC-Seq, and CUT&Tag for five histone modifications (HCR/LCR only), was carried out. See also Fig. S1a. (**d**) Jaccard similarity matrix of peak presence across epigenomic modalities, labeled to show similarity and diversity of function. (**e**) Example of a disrupted motif within differential peaks linked to *Acad11* expression. ATAC-Seq, H3K27ac, H3K4me1, and H3K4me3 peaks within the gene body of *Acad11* have a positive Pk2G link with *Acad11*. (**f**) Peak signal at the motif-disrupted region linked to *Acad11* (rolling mean bin size = 20, line = mean, shaded ± 95% CI). The nominal *p*-value from the DESeq analysis is provided for each modality. (**g**) Sequence comparison of HCR and LCR alleles within the Sp1 motif. The preferred Sp1 motif contains the G allele in HCR. 80% of the HCR population exhibit a G allele at this location, whereas 92% of the LCR population present the A allele. (**h-m**) HCR/LCR-F2 allelic concordance at *Acad11*. HCR/LCR relationship between (**h**) genotype and regulatory peak, (**i**) genotype and target gene, and (**j**) peak abundance and gene expression with Spearman’s correlation. F2 relationship between (**k**) genotype and regulatory peak with caQTL results printed, (**l**) genotype and target gene with eQTL results printed, and (**m**) peak abundance and gene expression, with regression line and Spearman’s correlation.

Approximately 90% of trait-associated common genetic variants are concentrated in noncoding regions of the genome^25^ where they can modulate transcription factor (TF) binding and gene expression through epigenomic activity^26^ in cell-specific and trait-relevant contexts. We hypothesized that selection-associated transcriptional divergence between HCR and LCR is reflected in line-specific epigenomic regulatory architecture shaped by noncoding genetic variation. Here we generate a comprehensive skeletal muscle epigenomic map of CRF using the HCR/LCR lines and the F2 cross, linking selected variants and their associated epigenomic architecture to the regulation of CRF-related gene expression.

## Results

### A noncoding variant at Acad11 links transcription factor binding to oxidative gene expression

To understand how noncoding genetic variation shapes epigenomic regulatory architecture of genes related to CRF, we profiled lateral gastrocnemius muscle from 128 HCR/LCR rats (Fig. S1a). We generated 546 multi-omic profiles across seven modalities including chromatin accessibility, five histone marks for chromatin state assessment, and gene expression (Fig. 1c-d, Fig. S2). Through integration of these data (Fig. S3, Fig. S4), we identified a distributed TF-centric enhancer architecture underlying the divergent transcriptome between HCR and LCR skeletal muscle. We then validated allelic effects on chromatin accessibility and gene expression in an independent HCR×LCR (F2) cross population^24^ (n=147) (Fig. 1c).

We illustrate this regulatory cascade at *Acad11*, a fatty acid dehydrogenase^27^, where a noncoding variant is predicted to increase Sp1 binding potential in parallel with increased chromatin accessibility and gene expression in HCR (Fig. 1e-g). A population-divergent variant at Sp1 motif position 10 (G allele frequency 0.80 in HCR vs 0.08 in LCR) strengthens potential binding of Sp1 in HCR (Fig. 1g). The HCR G allele coincides with neighboring E26 transformation-specific (ETS) family TF motifs within the region and elevated signal in ATAC-Seq, H3K27ac, H3K4me1, and H3K4me3 peaks (Fig. 1f) that are all positively linked to expression of *Acad11* (Fig. 1e). Concordant relationships between genotypes, peak abundance, and gene expression are observed across HCR/LCR and F2 studies (Fig. 1h-m). This regulatory logic identified at *Acad11* reflects a genome-wide pattern at mitochondrial metabolism genes, which we describe in detail below.

### Selection for running capacity drove regulatory divergence at distal skeletal muscle enhancers

We isolated the selection-associated CRF effects within the data by adjusting for sex, training, and acute exercise, and technical variation (Methods; Fig. S5, Fig. S6). In total, thousands of features significantly differed between HCR and LCR across all modalities and chromatin states (Fig. 2a, Fig. S7). On average, these HCR/LCR differential epigenomic regions reside 143.4 kb from the nearest TSS, consistent with widespread distal enhancer activity.

**Fig. 2:**
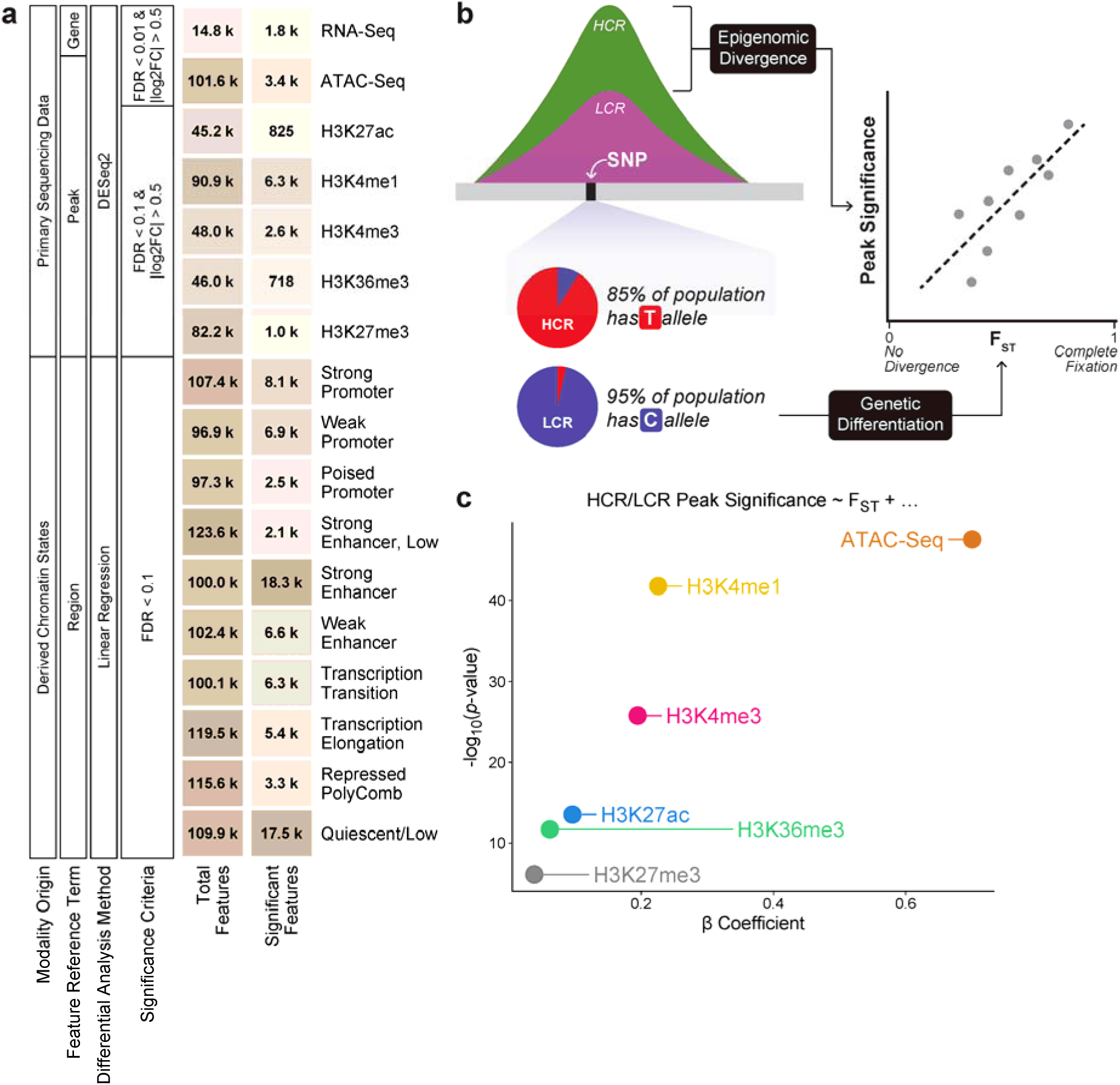
HCR/LCR epigenomic divergence colocalizes at genetically differentiated loci. (**a**) Summary of each data modality, including modality-specific terminology applied within this study, statistical thresholds selected based on statistical power, and the number of processed and significantly different HCR/LCR features. (**b**) Schematic showing comparison of population differentiation (F_ST_) and HCR/LCR epigenomic divergence carried out for each modality. (**c**) Association between F_ST_ and HCR/LCR peak significance from per-modality linear models shows that the most differential epigenomic peaks are also the most genetically divergent. The -log_10_(*p*-value) for the F_ST_ term in the model (y-axis) is plotted against the β coefficient (x-axis), with points colored by the modality.

Genes upregulated in HCR were predominantly enriched for mitochondrial-related processes including aerobic respiration, oxidative phosphorylation, fatty acid oxidation, and BCAA degradation (FDRs < 4.26×10^-9^) (Fig. S8a), consistent with previous studies^21,22^. A distinct signal related to angiogenesis was also identified (23 gene sets, highlighted in Fig. S8a). These pathways coincide with known CRF phenotypes of enhanced mitochondrial function^28^ and capillary density^29–32^ in HCR, mirroring that observed in humans with high intrinsic CRF^21,28,33–35^. These results provide a framework for interpreting how selection of distal regulatory elements shapes gene expression in skeletal muscle.

To test whether this widespread CRF-associated epigenomic divergence reflects underlying genetic selection for running capacity, we imputed genotypes from RNA-Seq derived single nucleotide polymorphisms (SNPs) (637,713 variants) and quantified allele frequency divergence between HCR and LCR by calculating per-variant fixation indices (F_ST_). Regressing HCR/LCR peak significance on peak-level F_ST_ while controlling for feature-specific artifacts (Methods; Fig. 2b) showed that genetic differentiation is more pronounced in differential epigenomic peaks, particularly for ATAC-Seq and H3K4me1 peaks (Fig. 2c). These findings indicate that selection for CRF associated regulatory variants is distributed across the genome, rather than at a single locus.

### Distal enhancer networks enriched for ETS and Mef2 motifs target oxidative and vascular genes

We linked regulatory elements to target genes by applying a peak-to-gene (Pk2G) correlation framework (Methods; Fig. 3a), annotating 88,440 peaks to 10,786 genes (FDR < 0.05, Fig. 3b). Peaks were linked to a median of 2 genes (Fig. 3c), reflecting the selectivity of enhancer-gene relationships^36,37^. For only 17.5% of linked peaks was the most strongly correlated gene the nearest gene (Fig. 3c), consistent with regulatory elements bypassing nearby genes to modulate distal targets^38–40^. The *Acad11* locus illustrates how multiple regulatory elements^41^ can correlate both positively and negatively with the expression of a single gene (Fig. 3d). This map defines a distributed regulatory architecture in which CRF-associated noncoding variation can cumulatively tune expression of metabolic and endothelial genes (Fig. S9).

**Fig. 3:**
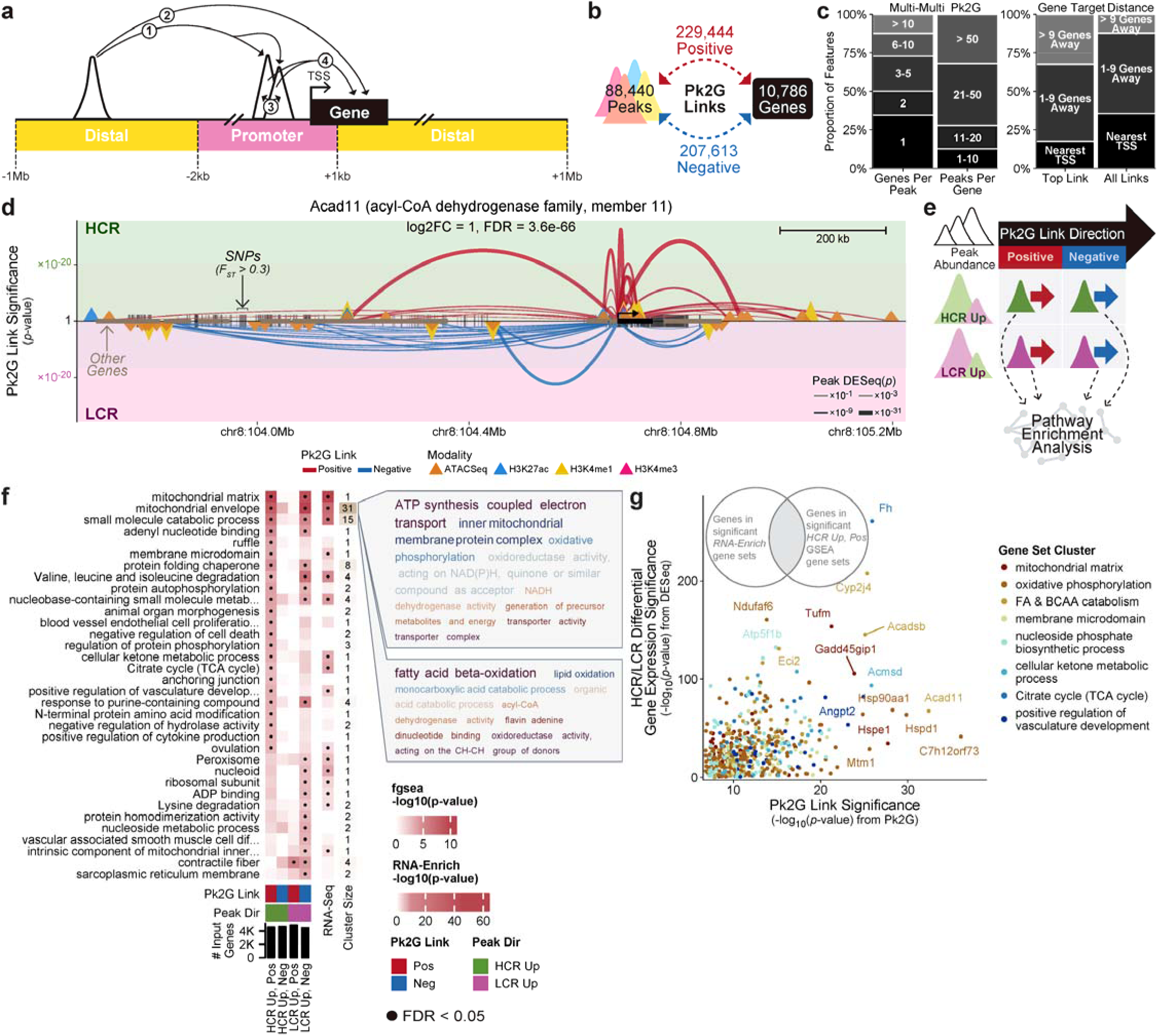
Distal peak-to-gene links connect CRF-associated enhancers to oxidative and vascular genes. (**a**) Correlation framework mapping epigenomic peaks (ATAC-Seq, H3K27ac, H3K4me1, and H3K4me3) to gene expression. Distal regulatory elements (yellow, TSS ± 1 Mb) are correlated with (1) promoter peaks (pink, TSS -2 kb, TSS +1 kb) and (2) gene expression. Promoter peaks are correlated with (3) one another and (4) gene expression. (**b**) Summary of the number of peaks and genes involved in the significant Pk2G links (FDR < 0.05) reflecting positive and negative peak-gene relationships. (**c**) Pk2G link distribution and target location. Proportion of features within binned categories of genes linked per peak and peaks annotated per gene (Multi-Multi Pk2G), and genomic distance from peaks to their linked gene targets, relative to the nearest TSS (Gene Target Distance). Shading intensity reflects bin order. (**d**) Pk2G links associated with a single gene, *Acad11* (log_2_FC = 1.04, FDR = 3.59×10^-66^). Regulatory elements (triangles, colored by modality) are positioned by genomic distance from the TSS (arrow) of the gene (black rectangle). Peaks with DESeq log_2_FC > 0 (HCR up) are located above the midline, and LCR up peaks are shown below. Arc height denotes the significance of the Pk2G links, colored by direction of Pk2G link, with linewidth proportional to DESeq significance of the regulatory element. Vertical ticks at the midline represent SNPs with F_ST_ > 0.3 and additional rectangles represent other genes within view. See additional examples in Fig. S9. (**e**) Schematic of four independent pathway enrichment analyses of activating (positive) and repressive (negative) regulatory relationships: 1) HCR up peaks positively linked to gene targets, 2) HCR up peaks negatively linked to gene targets, 3) LCR up peaks positively linked to gene targets, and 4) LCR up peaks negatively linked to gene targets. Genes were ranked by the Pk2G link significance. (**f**) Ranked gene set enrichment analysis of Pk2G targets linked to differential peaks for multiple analyses (columns), with the number of input genes indicated in the bottom bar plot. Significant gene sets (FDR < 0.05) are kappa-reduced. Representative terms are provided (‘•’ = significant) with annotated cluster sizes and RNA-Enrich results shown for comparison. The prominent gene sets collapsed into the two largest clusters are listed on the far-right side of the plot. (**g**) HCR-favored Pk2G mitochondrial metabolism genes compared to their differential expression. Comparison of the significance from the Pk2G correlation analysis included in the pathway enrichment analysis to the differential expression of the 537 genes that significantly enrich gene sets via GSEA of HCR Up, Pos gene links and RNA-Enrich differential gene expression. Y-axis shows -log_10_(*p*-value) from the HCR/LCR differential gene expression analysis, and the - log_10_(*p*-value) from the Pk2G link is plotted on the x-axis. Highly significant genes have strong relationships with epigenomic features, supporting coordinated regulation of the chromatin and gene expression.

Genes positively linked to HCR up peaks and negatively linked to LCR up peaks (Fig. 3e) were enriched for aerobic respiration, fatty acid and BCAA metabolism, as well as angiogenesis (Fig. 3f), mirroring RNA-Seq enrichment patterns (Fig. S8a). This agreement extended to individual genes, highlighting fatty acid and BCAA metabolism genes *Cyp2j4*, *Acadsb*, *Acad11*, *Mtm1*, and *Eci2* as leading candidates under selection-associated epigenomic regulation (Fig. 3g). Genes positively linked to LCR up peaks were enriched only for contractile fiber gene sets, highlighted by glycolytic type II fiber genes (Fig. 3f), consistent with the greater reliance on glycolytic metabolism during acute exercise observed in LCR^28^. When Pk2G correlations were recomputed within each line separately, the links driving these enrichments retained their direction and strength (Fig. S10a-b), indicating that selection reinforces the coordinated epigenomic regulation of energy metabolism and vascular development associated with high CRF.

Selection for running capacity was accompanied by divergence of the TF motif landscape within enhancers. We compared motif enrichment between HCR up and LCR up regions (Methods; Fig. 4a) and identified 146 differentially enriched motifs representing 78 TFs (Fig. 4b, Fig. S11a), largely independent of differences in TF expression (Fig. S12a). HCR up peaks positively linked to Pk2G targets were enriched for ETS-family TF motifs, whereas LCR up peaks negatively linked to Pk2G targets were enriched for Mef2 motifs (Fig. 4b, Fig. S11a). The corresponding target genes converged on mitochondrial oxidative phosphorylation, fatty acid oxidation, BCAA oxidation, mitochondrial ribosomes, and vascular development (Fig. S13a-b). The recurrence of these TF motifs across H3K4me1 enhancers linked to mitochondrial and vascular genes supports a model in which selection for running capacity shifts the regulatory architecture towards ETS-mediated transcriptional activation and away from Mef2-associated constraint (Fig. 4c, Fig. S14).

**Fig. 4:**
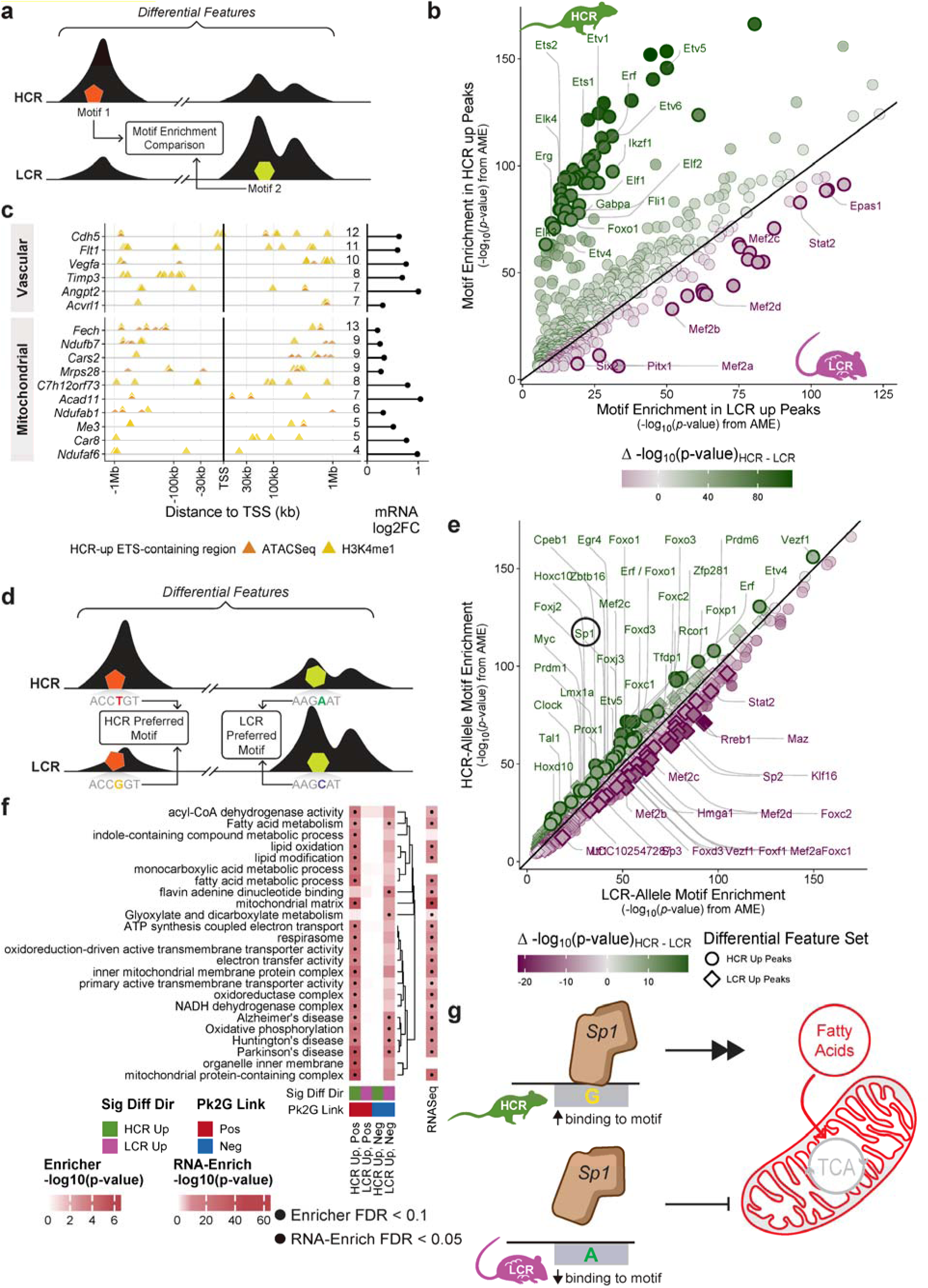
Divergent ETS and Mef2 motif landscapes distinguish HCR and LCR enhancers. (**a**) Schematic of motif enrichment comparisons between HCR up features with HCR alleles and LCR up features with LCR alleles, using the differential features defined in Fig. 2a. (**b**) Comparison of motif enrichment (analysis of motif enrichment (AME) -log_10_(*p*-values)) within H3K4me1 HCR up peaks (y-axis) and LCR up peaks (x-axis). Points further from the line of identity (slope = 1) and colored darker (green = HCR, magenta = LCR) represent motifs with strong differential enrichment, in which motifs within the top 10% Δ - log_10_(*p*-values) are outlined. See additional results in Fig. S11a. (**c**) Recurrent ETS-motif regulatory architecture at vascular and mitochondrial genes. Triangles represent peaks with ETS motifs with positive Pk2G links to genes in each row (color = modality), grouped by the gene set cluster that returned them (Fig. S13). The number of linked ETS-containing regions is printed on the right side of each row, followed by the HCR/LCR log_2_FC. See additional examples in Fig. S14. (**d**) Schematic of allele-specific differential motif enrichment analysis to detect TF binding differences due to line-divergent alleles within HCR up and LCR up regions. Motif enrichment is compared between HCR up regions with HCR alleles and HCR up regions with LCR alleles (left), as well as LCR up regions with LCR alleles and LCR up regions with HCR alleles (right). (**e**) Comparison of allele-specific motif enrichment (AME -log_10_(*p*-values)) between HCR alleles (y-axis) and LCR alleles (x-axis) within HCR up (circles) and LCR up (diamonds) H3K4me1 peaks. Points further from the line of identity (slope = 1) indicate HCR allele (green) and LCR allele (magenta) driven motif enrichment. Motifs within the top 10% Δ -log_10_(*p*-values) are outlined. See additional allele-specific results in Fig. S11b. (**f**) Over-representation analysis of genes linked to allele-disrupted motifs by direction (e.g., ‘HCR Up, Pos’ = HCR preferred motifs within HCR up peaks that are positively linked to the gene target, ‘•’ = FDR < 0.1 significance threshold) compared to RNA-Enrich (‘•’ = FDR < 0.05 significance threshold). (**g**) Visual summary demonstrating a model in which line-divergent alleles can alter TF binding and affect the expression of nuclear-encoded mitochondrial genes.

To determine whether specific line-divergent alleles contributed to this broader motif landscape, we compared HCR and LCR sequences containing variants with F_ST_ > 0.3 within HCR up and LCR up peaks (Methods; Fig. 4d). We identified 246 differentially enriched motifs representing 121 TFs (Fig. 4e, Fig. S11b), predominantly in the Cys2-His2 zinc finger (zf-C2H2), ETS, Forkhead, and basic helix-loop-helix (bHLH) TF families, all with established roles in skeletal muscle identity^42–48^ and mitochondrial function^49–55^. Importantly, 91.9% of the differentially enriched motifs correspond to TFs that are not differentially expressed between HCR and LCR (Fig. S12b), supporting a cis-regulatory contribution from divergent motif sequences rather than differences in TF transcript abundance. Fig. 1e-g illustrates one of 799 motif disruptions linking a line-divergent allele to gene expression through a predicted change in TF binding affinity (Methods). Genes positively linked to peaks containing HCR preferred motifs are enriched for fatty acid metabolism and aerobic respiration (FDR < 0.1; Fig. 4f), extending the pathway convergence observed across the broader motif landscape to the specific HCR/LCR allelic variation (Fig. 4g). These binding site-level disruptions accounted for only a subset of the motifs distinguishing HCR and LCR chromatin more broadly (Fig. S13). This suggests that noncoding variation propagates regulatory consequences beyond the immediately disrupted loci^56–58^.

### An independent F2 cross confirms allelic effects on chromatin accessibility and gene expression

Allele-dependent regulatory effects are preserved in an independent HCR×LCR (F2) cross population carrying the same variants^24^. We performed a large-scale validation study in the F2 population by generating RNA-Seq and ATAC-Seq libraries from gastrocnemius muscle, as well as genotyping the rats, to conduct expression quantitative trait locus (eQTL) and chromatin accessibility quantitative trait locus (caQTL) analyses (Fig. 1c). Accounting for linkage disequilibrium, we identified 4,747 genes with a significant eQTL (eGenes, FDR < 0.05) and 10,099 peaks with a significant caQTL (caPeaks, FDR < 0.05; Fig. 5a).

**Fig. 5:**
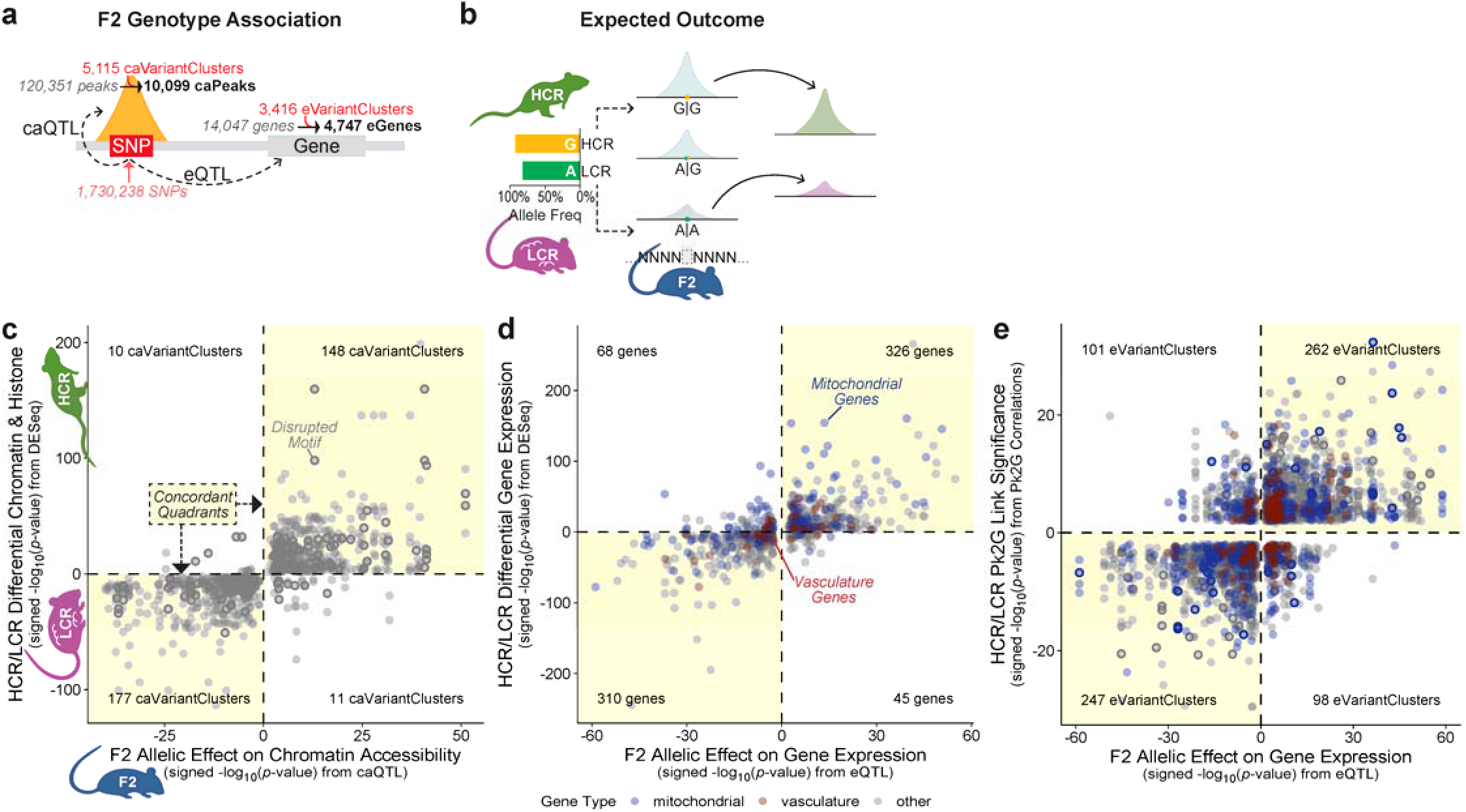
Allelic effects in an independent F2 cross recapitulate HCR/LCR epigenomic and transcriptomic divergence. (**a**) Overview of the validation study. F2 gene expression (RNA-Seq) and chromatin accessibility (ATAC-Seq) were related to genotypes via eQTL and caQTL analyses, respectively. (e/ca)VariantClusters = (e/ca)Variants with r^2^ ≥ 0.99 to the most significant variant. (**b**) Expected outcome. An HCR-predominant allele that positively associates with ATAC-Seq signal in F2 should correspond with elevated signal in HCR relative to LCR. (**c-e**) Concordance between F2 allelic effects (x-axis) and HCR/LCR differences (y-axis), restricted to variants with F_ST_ > 0.3. Shaded quadrants indicate directional concordance, with quadrant counts inset; concordance was tested using a binomial test (expected = 50%, alternative = “greater”). (**c**) F2 caQTL effects against HCR/LCR differential chromatin accessibility and histone modification abundance, 94% concordance (binomial *p*-value = 2.17×10^-63^). Outlined points indicate disrupted motifs by HCR/LCR allelic divergence. (**d**) F2 eQTL effects compared to HCR/LCR differential gene expression (DESeq FDR < 0.01), 85% concordance (binomial *p*-value = 1.63×10^-89^). Points are colored by the biological relevance of the gene (blue = mitochondrial, red = vasculature, gray = other). (**e**) F2 eQTL effects compared to HCR/LCR Pk2G links, 69% concordance (binomial *p*-value = 3.15×10^-17^). Points outlined as in c and colored as in d.

Genetic variants exert consistent regulatory effects across F2 and HCR/LCR populations (Fig. 5b), demonstrating that genetic divergence drives a substantial fraction of the regulatory differences between lines. We show significant directional concordance across three comparisons (binomial *p*-value ≤ 3.15×10^-17^ for all): F2 caQTL effects versus HCR/LCR epigenomic differences (94% concordance; Fig. 5c), F2 eQTL effects versus HCR/LCR gene expression differences (85% concordance; Fig. 5d), and F2 eQTL effects versus HCR/LCR Pk2G links (69% concordance, Fig. 5e). The Pk2G concordance further validates that the correlation-based links capture genuine regulatory relationships. These associations include genes involved in mitochondrial metabolism and vascular development (colored in Fig. 5d,e) and disrupted motifs from the HCR/LCR analyses (outlined in Fig. 5c,e). These results support that the broader epigenomic and transcriptomic differences observed between HCR and LCR reflect heritable regulatory divergence rather than line-specific artifacts.

Because a large portion of genetic variants associated with complex traits likely exert their effects through gene regulation^25,59^, the regulatory variants identified here should be of particular interest as entry points for dissecting the regulatory logic linking noncoding variation to CRF. Fig. 1e-m showcases one of the 293 validated variant-peak-gene examples that are likely to shape the regulatory landscape underlying CRF. The concordant allelic effects on chromatin accessibility and gene expression (Fig. 1h-i) are supported by significant association between the F2 genotypes at variant chr8:104691692 and chromatin accessibility of the F2 ATAC-Seq peak overlapping the HCR/LCR differential peak (caQTL β = 0.47, *p*-value = 5.1×10^-4^; Fig. 1k) and expression of *Acad11* (eQTL β = 1.11, *p*-value = 4×10^-37^; Fig. 1l). Additionally, the ATAC-Seq peak is positively correlated with *Acad11* expression in HCR and LCR (Fig. 1j) and F2 (Fig. 1m).

## Discussion

The skeletal muscle metabolic and vascular phenotypes that distinguish high from low CRF are well established^28–32^. What remains unclear is the molecular basis of the variability in these phenotypes within a heterogeneous population. Here, selection for CRF allowed us to stereotype the regulatory architecture of genes involved in mitochondrial oxidative metabolism and vascular development (Fig. 4, Fig. S13). By integrating transcriptomic and epigenomic profiles from N:NIH heterogeneous rat stock^18^ selectively bred for high and low running capacity^17^, we demonstrate that selection concentrated regulatory variants in a network of H3K4me1-marked distal enhancers linked to the expression of hundreds of genes central to CRF. Concordant allelic effects on chromatin accessibility and gene expression in the independent F2 cross population confirmed regulatory effects established in the HCR/LCR study (Fig. 1h-m, Fig. 5). These CRF-associated epigenomic regions are characterized by high densities of ETS-family TF motifs in HCR and Mef2 TF motifs in LCR, suggesting that selection acted on noncoding variants distributed across many enhancers that cumulatively^60–62^ shape divergent transcriptional profiles, as proposed for complex trait architecture^63^.

Multiple regulatory elements converge on the same target genes (Fig. 3d, Fig. S9, Fig. 4c, Fig. S14) with an architecture reminiscent of the ‘shadow’ enhancers described for developmental loci^64^. Distributing control across several enhancers per gene buffers expression against mutations that may disrupt any individual enhancer^65^. Such redundancy may also support context-specific tuning^66^, allowing relative contribution of each enhancer to vary across conditions^67,68^, and in some cases, across allelic background or individuals^62,63,69^, a property that remains underemphasized in complex trait genetics.

Our findings extend the prevailing model of mitochondrial gene regulation in skeletal muscle, which centers on PGC-1α coactivation of Nrf1 and Nrf2/Gabpa at proximal promoters^70,71^, to include distal enhancer networks shaped by CRF-associated noncoding variation. Expression of these TFs and cofactors differs only modestly between lines (*Ppargc1a* log_2_FC = 0.49, FDR = 1×10^-14^; *Nrf1* log_2_FC = -0.03, FDR = 0.23; *Gabpa* log_2_FC = - 0.14, FDR = 3.5×10^-11^). Such shifts in broadly acting regulators cannot account for gene-specific divergence observed across thousands of distal enhancers linked to oxidative metabolism and angiogenesis targets (Fig. 3, Fig. 4a-c, Fig. S13, Fig. S14), which is maintained in the segregating F2 population (Fig. 5). Notably, *Gabpa* expression is slightly lower in HCR even though ETS motifs are enriched in HCR up peaks (Fig. 4b). We identify this complementary cis-regulatory layer, a network of distal ETS-family enriched enhancers (including Gabpa motifs; Fig. 4b), in which divergent TF binding opportunity prioritizes which metabolic genes are able to respond to PGC-1α mediated transcriptional enhancement in a given regulatory context.

Selection for CRF acted less through the abundance of canonical mitochondrial regulators than through the accumulation of noncoding variants that reweigh distal enhancer motif composition from Mef2 toward ETS-family TFs. ETS-enriched enhancers were positively linked to oxidative metabolism genes in HCR, while Mef2-enriched enhancers were negatively linked to these genes in LCR (Fig. 4b-c, Fig. S13). Direct allelic disruption of TF motifs accounted for only part of this shift (Fig. 4e-f), in which the remaining divergence likely reflects proximal and distal variant effects on TF binding, chromatin accessibility, and enhancer activity^72–77^. This ETS-associated shift extends to angiogenesis genes (Fig. S13b), suggesting that selection targeted a shared regulatory node coupling metabolic and vascular programs. This ETS-dominant regulation in HCR is reflected in phenotypes of greater skeletal muscle mass^30^, increased skeletal muscle mitochondria^78–80^ and capillary density^29–32^, and higher type I and IIa fiber content^29,81^ relative to LCR, supporting metabolic efficiency and delayed depletion of muscle glycogen during intense exercise^28^.

This co-regulation may have a developmental basis. Skeletal muscle and endothelial cells derive from common bipotent progenitors in which ETS-family TFs, including Fli1 and Gabpa, play lineage-determining roles^57,82–84^. The prominence of CRF-divergent H3K4me1-marked enhancers raises the possibility that some of these regulatory elements were established during this shared developmental window and later co-opted for tissue-specific gene regulation. This could explain how selection on ETS binding potential coordinately elevates oxidative capacity and vascular supply in HCR.

Coordinately raising oxidative capacity at multiple entry points of mitochondrial electron flux through ETS-enriched enhancers linked to fatty acid oxidation, TCA cycle, and oxidative phosphorylation genes (Fig. 6) broadens prior reports of ETS-family TFs binding at individual mitochondrial gene promoters^49,85^ to a distributed distal enhancer network. Acyl-CoA dehydrogenases are prominent targets, consistent with the elevated capacity for β-oxidation in HCR skeletal muscle. This is achieved by direct routing of electrons to coenzyme Q (CoQ) via the electron-transfer flavoprotein (ETF) and ETF-QO (*Etfdh*, log_2_FC = 0.54, FDR = 5.5×10^-83^)^86^. In conjunction with epigenomic regulation of acyl-CoA dehydrogenase isoforms that mitigate oxidative damage through peroxidized fatty acid metabolism^87,88^, this electron rerouting can protect against the accumulation of lipotoxic intermediates^89^, short-chain acyl-CoA^90^, and reactive oxygen species^91^ that characterizes mitochondrial dysfunction in type 2 diabetes^92–94^. Notably, many of the fatty acid and BCAA-degrading dehydrogenases upregulated in HCR are evolutionarily ancient genes derived from α-proteobacteria with orthologs across Archaea, Bacteria, and Eukaryota^95,96^, suggesting that while their coding sequences and enzymatic function have been conserved across billions of years, regulatory adaptation continues to adjust their expression based on energy availability and demand.

**Fig. 6:**
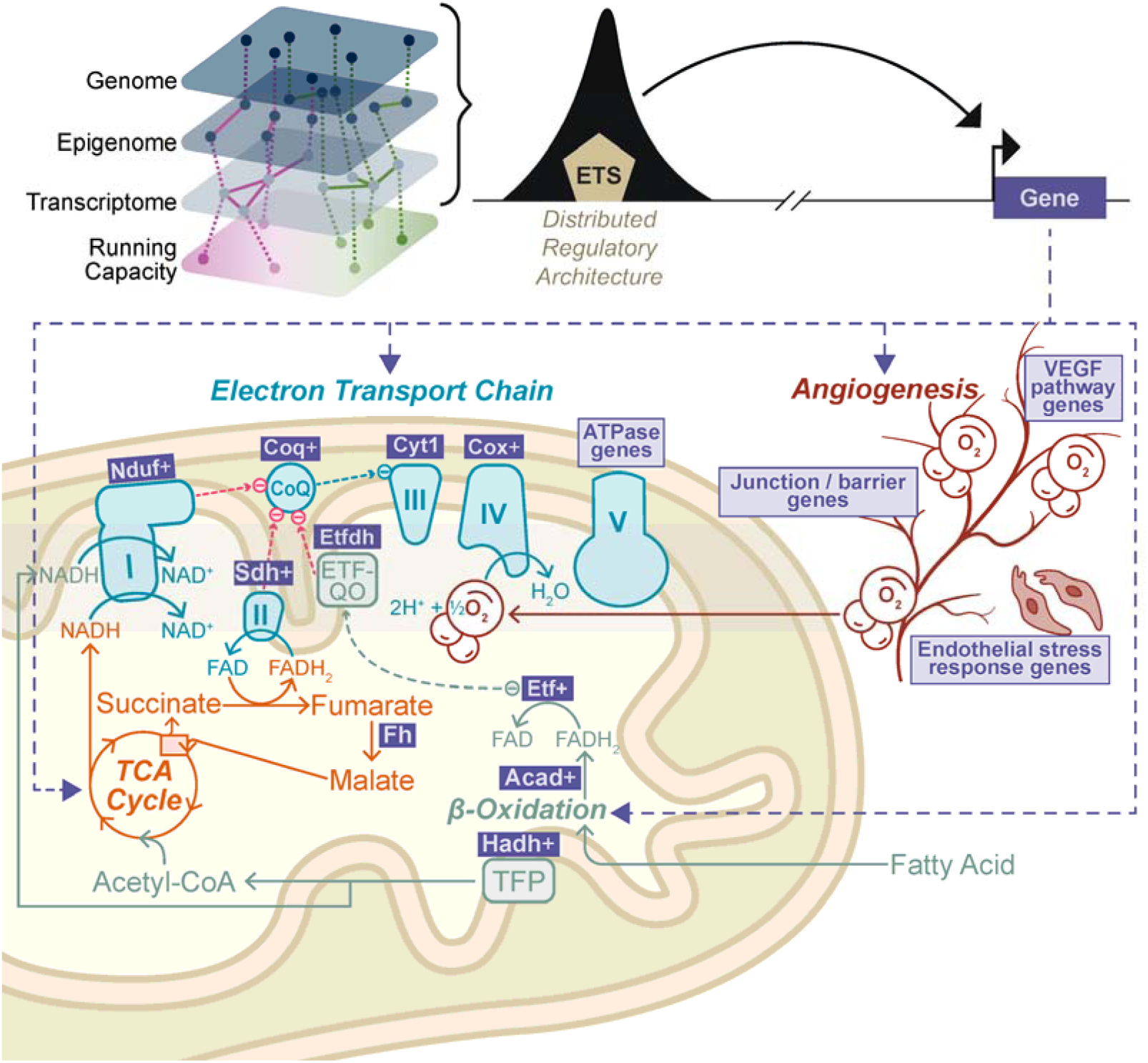
ETS-mediated regulation of mitochondrial energy metabolism supports oxidative efficiency in high CRF muscle. Model of ETS TF motifs within enhancer regions linked to upregulated genes involved in fatty acid oxidation (acyl-CoA dehydrogenases, *Hadh*, *Etfa*/*Etfb*, *Etfdh*), TCA flux (*Fh*: log_2_FC = 0.68, FDR = 1.47×10^-257^, rank = 3), electron transport (subunits of complexes I-V), and endothelial pathways, which together support efficient oxidative metabolism in HCR skeletal muscle. A gene symbol followed by ‘+’ indicates that multiple family members are targeted. TFP = trifunctional protein.

Several limitations frame these results. The HCR/LCR comparisons are associative by design. We address this with an independent F2 cross, where eQTL and caQTL mapping confirms that line-divergent alleles act in cis on chromatin accessibility and gene expression (Fig. 5c-e), though we do not directly perturb individual enhancers to establish causality at single loci. Our profiling is restricted to bulk lateral gastrocnemius, so differences in cell-type composition between lines may contribute to some signals. The extent to which this regulatory architecture is conserved in human skeletal muscle is directly addressable with existing multi-omic cohort data and is the subject of ongoing work.

Intrinsic CRF is a complex trait, a determinant of cardiometabolic health, and a reproducible predictor of longevity^19^. Selective breeding of rats for running capacity amplified phenotypes relevant to human cardiometabolic health^20^, with the greatest impact on skeletal muscle as observed by fewer mitochondrial gene expression differences in heart^23^, liver^97^, and adipose tissue^98^ between HCR and LCR lines. The regulatory variants identified here mark candidate genomic loci for understanding how noncoding variation shapes oxidative capacity in human skeletal muscle, a trait tied to sarcopenia^99^, aging^100^, obesity, and insulin resistance^101,102^. Beyond CRF, our framework consisting of multi-omic profiling and integration to identify peak-gene regulatory networks, nomination of TFs through motif enrichment, and cross-population validation of allelic effects provides a template for dissecting complex trait architecture (Fig. S3). Together, these results link polygenic selection on a heritable cardiometabolic trait to specific cis-regulatory variants and the oxidative metabolism transcriptional programs they control in the skeletal muscle.

## Methods

### HCR/LCR Study Design

A total of 128 rats from generation 39 of the HCR/LCR lines (38 weeks of age) were included in this study, including equal numbers of females and males (Fig. S1a). Animal procedures were approved by University of Michigan IACUC, Protocol PRO00006206. Half of the rats, deemed the “training” group, underwent a six-week exercise regimen consisting of three days per week (Monday, Wednesday, and Friday) of 30 minutes of supervised treadmill running up a 15-degree incline. The treadmill speed for each week of training was set to 75% of the average maximal running speed determined from a test performed the week prior. This weekly maximal running test was performed separately on a group of 3-4 rats from each line (LCR and HCR) and sex using the standard LCR/HCR test protocol, as described previously^17^. When performed, this test took the place of the normal training procedure for that group of rats on Fridays. The other half of the rats (deemed the “sedentary” group) were subjected to once-weekly treadmill acclimatization on Tuesdays for six weeks. This consisted of 5 minutes of treadmill walking at a slow pace (5 m/min) and 15-degree incline, followed by an additional 25 minutes of resting on a motionless treadmill, before being returned to their home cages. The training or acclimatization process was stagger-started over the course of three weeks to allow the final bout of exercise with dissection and sample collection to be performed after a consistent 72-96-hour washout period following the last bout of training.

Once training was complete and the washout period had passed, rats from both the “trained” and “sedentary” groups were subjected to an acute ascending-effort exercise bout using the standard LCR/HCR test protocol, with variation of the endpoint to allow sample collection at four different timepoints relative to initiation of exercise. The “rest” timepoint was collected immediately after animals were removed from their home cage. These rats did not perform any acute exercise on the day of sample collection. “Half-max” samples were collected from a group of rats that underwent the test protocol but were removed from the treadmill at half of their predicted maximum running time, which was determined by the group-specific averages measured on the last Friday of the training period, as already described. The “half-max” timepoints were as follows: HCR females, 40 minutes; HCR males, 35 minutes; LCR females and males, 10 minutes. Samples collected at “maximum” exercise were collected from rats run to exhaustion according to the standard LCR/HCR test protocol. Samples collected during “recovery” were collected from rats that were run to exhaustion using the standard test protocol, then returned to their cage for 1 hour prior to dissection. The running time and distance achieved by animals in the “maximal” and “half-max” groups were recorded for each animal. At the time designated for dissection, which was immediately after removal from the treadmill for the “half-max” and “max” groups, and as described above for “rest” and “recovery” groups, rats were placed in an anesthesia induction chamber ready with isoflurane anesthetic vapor. Once anesthesia was induced (∼15 seconds) the animal was transferred to a nose-cone connected to a metered anesthetic-delivery machine and was maintained under deep anesthesia during sample collection.

### Accounting for Experimental Variability in the HCR/LCR Study

This study is part of a larger effort to investigate transcriptomic and epigenomic changes in skeletal muscle as a response to exercise training and an acute exercise bout in relation to CRF, but the presented results focus on the intrinsic differences between HCR and LCR. While the six-week training regimen resulted in increased running capacity in both HCR and LCR (Fig. S1d), there were minimal transcriptomic and epigenomic differences 72-96 hours after the last exercise bout (data not shown). However, an acute exercise bout induced considerable transcriptomic and epigenomic changes (data not shown). In this study, we included all samples, when possible, to maximize our power and carried out extensive QC on count data associated with RNA-Seq, ATAC-Seq, and CUT&Tag modalities. We normalized for technical and experimental variables (acute exercise, training group, and sex) to isolate the HCR/LCR effect, as described in the applicable methods sections. A detailed analysis of exercise-induced experimental contrasts will be presented in future work.

### Preparation of High-Throughput Sequencing Libraries

#### CUT&Tag Nuclei Isolation and Library Generation

Skeletal muscle nuclei were isolated from the gastrocnemius muscle of 60 HCR/LCR rats following the bulk nuclei isolation protocol published in Orchard et al.^103^. When isolating nuclei prior to CUT&Tag library preparation, modifications were made at the following steps: 3) a rotating plate was used to rock the samples for 3 minutes, 11 & 12) the nuclei pellet was spun at 500 × g, 13) a final concentration of 5 µM DAPI was used to count the nuclei. Throughout the protocol, CUT&Tag wash buffer was used to resuspend the sample and pre-wet the filters.

We profiled five histone modifications via CUT&Tag^104^ following the “CUT&Tag-direct with CUTAC V.3” protocol^105^. To reduce batch effect across experimental contrasts, samples were randomly assigned to five batches to carry out 360 CUT&Tag reactions at the bench (Fig. S1b). All six reactions for a given sample were processed in the same batch. In the case of library preparation failure, libraries were re-prepared in a later batch. A sixth preparation batch including four libraries was added to complete the library preparation. From the nuclei isolation step, 15K nuclei were prepared for each of the six CUT&Tag reactions per sample. Primary antibodies for histone modifications probed include H3K27me3 diluted 1:100 (Cell Signaling, 9733), H3K27ac diluted 1:100 (Abcam, ab4729), H3K4me1 diluted 1:100 (Cell Signaling, 5326), H3K4me3 diluted 1:100 (Diagenode, C15410003-50), and H3K36me3 diluted 1:100 (Active Motif, 61101). A negative control where no antibody was added was included for each sample. Bioanalyzer profiles were assessed to confirm success of experiment (absence of DNA in the no-antibody control), confirm nucleosomal patterns of histone modification samples, and measure DNA concentration for equimolar pooling of libraries to sequence. The 300 libraries where a primary antibody was included were sequenced. Libraries were re-randomized for sequencing batches, referred to as “flowcell” for the purposes of this study. Libraries were sequenced using paired-end, 150 bp reads on the NovaSeq platform at the University of Michigan Advanced Genomics Core to an average depth of 7.7 M paired-end reads per library.

Five CUT&Tag libraries from a single sample were excluded because that sample was a consistent outlier in PCA across all histone modifications. At the time of experiment, the muscle tissue also appeared discolored with an unusually powdery consistency. As a result, 59 samples (295 libraries) were included in our final CUT&Tag dataset.

#### ATAC-Seq Nuclei Isolation and Library Generation

Flash frozen gastrocnemius muscle samples from 128 HCR/LCR rats and 149 F2 rats were processed independently. In the HCR/LCR preparation, the samples were divided into approximately 21 batches comprising six samples each representing each of the six conditions, which were then re-randomized across sequencing batches (Fig. S1c). F2 samples were divided into 47 batches, with six samples in each batch. For each of the samples, tissue pieces weighing 50 mg were cut on dry ice using a frozen scalpel to prevent thawing. The samples were pulverized using liquid nitrogen and a cell crusher. Within each batch, each of the six samples was resuspended in 1 mL of ice-cold PBS to perform six different nuclei isolations. We developed a customized protocol (Supplemental Protocol S1) from the previously published ENCODE protocol, and used it to isolate nuclei, which is compatible with bulk ATAC-Seq. The nuclei were counted on a Countess II FL Automated Cell Counter, and the appropriate volume of the suspension for 50K nuclei was spun down and used for the downstream transposition reaction (ENCODE Project Consortium 2012, 2012). The 50K nuclei were transposed, PCR amplified, and the libraries were generated in duplicate by adding appropriate sample indexes. Five HCR/LCR and eight F2 ATAC-Seq samples were excluded due to failure in pre-sequencing QC or unintentional duplication, resulting in 123 samples in the HCR/LCR and 141 samples in the F2 ATAC-Seq datasets. Libraries were sequenced using paired-end, 51 bp reads on the NovaSeq platform to an average depth of 63M paired-end reads per HCR/LCR sample (across 2-4 read groups) and 67M paired-end reads per F2 sample (across 1-4 read groups).

#### RNA-Seq Library Generation

Frozen gastrocnemius muscle from 128 HCR/LCR and 150 F2 samples were pulverized in liquid nitrogen. RNA was subsequently extracted using the Zymo Direct-zol kit. Stranded RNA-Seq libraries were generated using poly(A) selection and were sequenced using paired-end, 150 bp reads on the HiSeq platform to an average depth of 39M paired-end reads per HCR/LCR sample and 37 M paired-end reads per F2 sample.

### Data Analysis

All analyses were performed using R Statistical Software (v4.3.2; 2023-10-31) and Python (v3.10.4). Commonly used R packages include tidyverse^106^, GenomicRanges^107^, plyranges^108^, and ComplexHeatmap^109,110^. Snakemake^111^ was used to create pipelines used throughout the analysis. Sequencing data were aligned to mRatBN7.2 (rn7, Ensembl release 109^112^).

### Processing of RNA-Seq Data

Fastq files were trimmed using the TrimmomaticPE function (Trimmomatic^113^ v0.36) with the following parameters: ILLUMINACLIP:TruSeq3-PE.fa:2:30:10:2:keepBothReads SLIDINGWINDOW:5:20. Reads were aligned using STAR^114^ (v2.7.6a-aekjdpr) with the parameters “-c --outSAMtype BAM SortedByCoordinate”. To create the count matrix, gene counts from QC.geneCounts.txt.gz files produced by QoRTs^115^ command QC with parameter –stranded and the same GTF file used in the alignment were used, returning 30,454 genes. The count matrix was filtered to remove non-autosomal genes and genes where all samples had 0 counts. For the HCR/LCR analysis, RNA-Seq counts were further filtered to include genes where more than 10 samples had a count value > 5, resulting in 14,781 genes in the dataset used in this analysis. VST-normalized counts, adjusted for experimental contrasts and technical variables, were used for visualization.

### Processing of ATAC-Seq and CUT&Tag Data

Fastq file adapters were trimmed using cta version 0.1.2 (https://github.com/ParkerLab/cta) and then aligned to the rn7 genome (Ensembl file Rattus_norvegicus.mRatBN7.2.dna.toplevel.fa.gz) using the bwa^116^ *mem* version 0.7.15 with parameters “-I 200,200,5000” and then sorted using samtools^117^ version 1.13. The bwa “-M” flag was used for aligning ATAC-Seq data only. ATAC-Seq libraries were prepared with multiple read groups, which were merged using samtools^117^. Duplicates were marked using picard^118^ *MarkDuplicates* version 2.25.5 with parameters “-ASSUME_SORTED true -VALIDATION_STRINGENCY LENIENT”. Properly paired and mapped autosomal reads with high quality alignments (≥ 30) were retained using samtools^117^ *view* with parameters “-h -f 3 -F 4 -F 8 -F 256 -F 1024 -F 2048 -q 30”. Bedtools^119^ version 2.30.0 was used to remove the custom blacklist regions created specifically for this data. BAM files were converted to BED files and peaks were called using the *callpeak* command from MACS2^120^ version 2.2.4. Narrow peaks were called for

H3K27ac, H3K4me1, and H3K4me3. Broad peaks were called for H3K27me3 and H3K36me3. Narrow and broad peaks were called for ATAC-Seq. Parameters “-g hs -B --keep-dup all -q 0.05” were included for all peak calls. H3K27me3 and H3K36me3 peak calling included the “--broad” flag. ATAC-Seq broad peaks had the added parameters, “--SPMR -f BED --nomodel --shift -100 --seed 762873 --extsize 200 --broad”. ATAC-Seq narrow peak call had the added parameters, “--SPMR -f BED --nomodel --shift -37 --seed 762873 --extsize 73 - -call-summits”. Ataqv^121^ was used to generate QC metrics used for further data normalization. For visualization, raw signal was smoothed on a per-sample basis by applying a rolling mean.

Data quality and reproducibility were assessed at an individual and global level. We show the epigenomic signal from an individual sample surrounding *Gapdh*, a housekeeping gene expressed in skeletal muscle (Fig. S2a). H3K4me3, ATAC-Seq, and H3K27ac signals are observed at the transcription start site (TSS), indicative of an open and active promoter region. Moderate levels of H3K36me3 signal, along with low levels of the repressive H3K27me3, reflect active transcription of the gene. Study-wide consistency of our data is exemplified by signal concordance at representative loci across modalities (Fig. S2b) and within-modality clustering from principal component analysis (PCA) of genome-wide fragment counts (Fig. S2c). Notably, PC1 exhibited separation between repressive and active marks, while PC2 separated different classes of active marks, demonstrating the reproducibility and biological specificity of the diverse epigenomic profiles.

### ATAC-Seq and CUT&Tag Count Matrix Construction for HCR/LCR Study

MACS2 called narrow peaks (H3K27ac, H3K4me1, H3K4me3), broad peaks (H3K27me3, H3K36me3), and narrow summits extended ± 150 bp (ATAC-Seq) were used to create the consensus peak matrices. For each unique line + sex + acute exercise group, peaks were merged and the number of distinct samples that the peak occurred in was tallied. Peaks were retained if they were present in the majority of samples for each line + sex + acute exercise group (Fig. S15). Peaks present in the majority of any of the 12-16 unique groups (16 for ATAC-Seq; 12 for CUT&Tag, which lacks half-max) were merged to create the master peak matrix. In R, plyranges^108^ *reduce_ranges* was used to create the master peak set for each modality and chromVar^122^ *getCounts* with arguments “paired = FALSE, by_rg = FALSE, format = ‘bam’” was used to obtain counts within the consensus peak regions from the pruned BAM file. VST-normalized counts were used for visualization. Normalized counts were adjusted for sex, exercise group, and technical variables (preparation batch, sequencing batch, latent variables) while protecting for line.

### Custom Blacklist Creation for rn7 Genome

A widely used blacklist for the rn7 genome was not available at the time of analysis, so a blacklist custom to this dataset was created following the process outlined in Amemiya, 2019^123^. A combination of 336 CUT&Tag and ATAC-Seq libraries prepared from 56 rats were used to create this blacklist region file. Briefly, 1 kb bins with 100 bp overlapping regions were created using bedtools^119^ version 2.30.0 *makewindows* and the number of reads present in each bin were counted. To detect erroneous signal, quantile normalization (broman::normalize^124^) was carried out on binned count matrices, for each data modality independently. A region was included in the final blacklist based on the presence of outlier signal in all six data modalities. To consider the fact that some datasets contain sequencing information for differing areas of the genome, the region was also included in the blacklist file if the median value of the quantile normalized matrix was 0. This resulted in 278 blacklist regions with cumulative width of 848 kb.

### Latent Variable Discovery for ATAC-Seq, CUT&Tag, and RNA-Seq

RUVSeq^125^ was used to generate and trial 1-10 latent variables. Control peaks consisted of features with a DESeq *p*-value > 0.1 across all known experimental and technical contrasts, with the number of control peaks used ranging from 2,000-14,000+. Latent variable inclusion was determined by their ability to improve biological discovery assessed by the number of significant DESeq features (Fig. S6a) and observed variability in PCA when added to the starting model (Fig. S6c-d), while ensuring that the latent variable(s) show a relationship with the ataqv^121^ QC attributes and minimal relationship with the binarized experimental variables (Fig. S5a-g). When multiple models yielded similar results, the simpler model was preferred.

### RNA-Seq, ATAC-Seq, and CUT&Tag Differential Analysis for HCR/LCR study

DESeq2^126^ was used to perform differential analysis on CUT&Tag histone modification peaks (H3K27ac, H3K4me1, H3K4me3, H3K27me3, H3K36me3), ATAC-Seq peaks, and RNA-Seq gene counts using the model specified in Fig. S6b. Technical variables, experimental contrasts, and latent variables were included in the DESeq model. For the ATAC-Seq and RNA-Seq analyses, a stringent significance threshold of FDR < 0.01 and |log_2_FC| > 0.5 was applied. Given the smaller sample size in the CUT&Tag libraries, a more lenient cutoff of FDR < 0.1 and |log_2_FC| > 0.5 was utilized for histone modifications. A log_2_FC threshold was included for CUT&Tag to compensate for the more lenient FDR, and for other DESeq-based analyses to maintain consistency.

### Chromatin State Discovery

ChromHMM^127^ was used to call chromatin states in 200 bp bins across the genome. Custom rn7 files were created for chromosome sizes, coordinates, and anchor files using the R packages GenomicFeatures^107^ and TxDb.Rnorvegicus.UCSC.rn7.refGene.

The 56 samples with complete ATAC-Seq, H3K27ac, H3K4me1, H3K4me3, H3K36me3, and H3K27me3 modalities (336 total libraries) were integrated to discover chromatin states. Custom blacklist regions were removed from BAM files, which were then down sampled to the lowest read depth (> 2M) per modality. If the lowest read depth was < 2M reads within a modality, then BAM files with read depths > 2M were down sampled to 2M and BAM files with read depths < 2M were not down sampled. There were three libraries with read depths ranging from 1.5-1.7M. ChromHMM state models ranging from 2-20 were assessed to choose the 12-state model (Fig. S4a), which presented minimal change in the log likelihood of the subsequent model and was most informative, given the combination of epigenomic marks. Emission states were manually assigned the following chromatin state names (with associated abbreviations): E1 = Strong Enhancer, Low (EnhS_E1), E3 = Weak Enhancer (EnhW_E3), E4 = Strong Enhancer (EnhS_E4), E5 = Weak Promoter (PromWk_E5), E6 = Transcription Transition (TxTrans_E6), E7 = Strong Promoter (PromA_E7), E8 = Inactive/Poised Promoter (PromPois_E8), E10 = Repressed PolyComb (ReprPC_E10), E12 = Transcription Elongation (TxnElon_E12). For example, we observe greater likelihood of a Strong Promoter chromatin state call, in comparison to other states, at promoter regions of genes (Fig. S4b). Emission states E2, E9, and E11, returned states associated with low signal, were assigned the “Quiescent/Low” state and summed together into a single “Quiescent/Low”, labeled as QuiesSum_EX in the results data tables.

### Chromatin State Differential Analysis

ChromHMM-derived posterior probabilities were prepared for group analysis by retaining 200 bp regions where any line, sex, or acute exercise group called a non-quiescent state, based on the highest initial group averaged state probability. In the case of a tie, a non-quiescent state assignment was prioritized. This was followed by summing quiescent states into one “QuiesSum” state. This produced posterior probability matrices that included 1,160,951 200 bp regions. For each chromatin state, 200 bp regions where the sum of posterior probabilities across samples was greater than 0 were included for differential analysis. The count of merged regions included in this analysis is provided in Fig. 2a. Linear regression was applied to posterior probabilities that were inverse rank normalized (ties = “random”) across samples, using the model ∼ *Line* + *Sex* + *Acute Exercise Group*. *P*-values for the HCR/LCR line contrast were obtained and FDR-adjusted. We excluded emission state E1 (labeled as “Strong Enhancer, Low”) from further analyses because it showed consistently low ChromHMM posterior probabilities across samples. At most, 22/56 samples had a posterior > 0.1 for this state, and its 99^th^ percentile posterior was 0.07. In contrast, the other states had many 200 bp regions where all 56 samples returned a posterior > 0.1, with the 99^th^ percentiles of posteriors being 0.65 for Repressed PolyComb, 0.83 for Transcription Transition, and > 0.96 for the remaining 7 chromatin states. Because E1 posteriors were near-zero in most samples, a group-level call, which is based on the averaged posteriors from samples within the group, was unlikely. Chromatin state regions with an FDR < 0.1 were considered significantly different due to a smaller sample size. A magnitude of change threshold was not applied to ChromHMM-derived states due to the inverse normal transformation applied to posterior probabilities used in the linear model.

Additional QC of ChromHMM assignment of state probabilities across genome-wide loci to enable systematic quantification of regulatory divergence consisted of viewing the epigenomic signal alongside ChromHMM posteriors. A representative chr14 locus illustrates HCR/LCR chromatin state divergence where increased ATAC-Seq and H3K4me1 signal in HCR support a Strong Enhancer state, whereas LCR exhibit a Quiescent/Low state (Fig. S4c).

### Pathway Enrichment Analysis

To ensure consistency across transcriptomic and epigenomic comparisons, a custom list of gene sets to probe was created by retrieving gene sets and their associated genes (Ensembl IDs) from GOBP, GOMF, GOCC, and KEGG via msigdbr^128^. Missing Ensembl or Entrez ID matches were filled in where possible using the *biomaRt*^129^ package to retrieve gene information from the *rnorvegicus_gene_ensembl* v109^112^ database, ensuring consistency in gene set annotations. Categories with 10 to 500 genes were probed for pathway analysis. Resulting *p*-values were FDR adjusted for each individual database using the Benjamini-Hochberg method^130^. Unless stated otherwise, categories returning an FDR < 0.05 were considered significant.

Pathway enrichment analysis was performed independently on all data modalities. RNA-Enrich^131^ was used for pathway enrichment analysis of differential gene expression data which included all genes, ranked by *p*-value with direction of change provided.

To assess the processes that Pk2G link target genes were involved in, HCR/LCR differential peaks passing the FDR threshold provided in Fig. 2a were queried. The list of input genes was separated by direction of peak change and direction of relationship between the peak and target genes. The number of input genes for each analysis is as follows: HCR up peaks positively linked to 4,600 genes, HCR up peaks negatively linked to 4,678 genes, LCR up peaks positively linked to 4,889 genes, and LCR up peaks negatively linked to 4,490 genes. The fgsea function from the fgsea^132^ R package was used to carry out pathway enrichment on these sets of genes, ranked by significance of correlation.

The enricher function from the ClusterProfiler^133^ R package was used to carry out over-representation analysis using specified target genes and all genes expressed in the skeletal muscle as the background list (n = 14,781). When the target genes of regions contributing to allele-specific differential motif enrichment analysis were considered, the HCR up pos analysis used 330 genes, HCR up neg analysis used 279 genes, LCR up pos analysis used 272 genes, and LCR up neg analysis used 281 genes as the target list.

To reduce redundancy and clarify comparisons across independent analyses, the clustering approach described by Metascape^134^ was applied. Functions created to carry out gene set reduction by kappa clustering are published on GitHub (https://github.com/weitzela/reducekappa, commit 04eb74d). Briefly, pairwise kappa values were calculated to capture the similarity of genes associated with all categories across (one or multiple) analyses. Kappa scores were min/max normalized to adjust for negative values. The kappa matrix was converted to a distance matrix by subtracting the kappa scores from 1, which was then used for hierarchical clustering (method = “average”). Clusters were identified using a threshold height of 0.7. The representative category for each cluster was selected as the term with the lowest *p*-value across all analyses. For comparisons across multiple modalities, values associated with the most significant category within each cluster were retained for individual analyses and plotted, even if that category was not named the “representative” one.

### Correlation of Distal Regions with Promoter Regions and Gene Expression

To expand annotation links between epigenomic features and genes, we conducted a correlation analysis utilizing RNA-Seq, ATAC-Seq, H3K27ac, H3K4me1, and H3K4me3 data modalities. We extended the approach described in Currin et al.^135^, detailed further here. This adaptable peak-gene annotation method is made publicly available as an R package (https://github.com/weitzela/PeakGeneNet).

We used the limma::removeBatchEffect^136^ function to adjust the DESeq VST counts (blind = FALSE) for technical variables such as prep date, flowcell, and latent variables discovered/used for differential analysis. To ensure that the line effect remained when evaluating the relationship between peaks and genes, non-*Line* experimental contrasts were further adjusted for via linear regression. Promoter regions were defined as regions 2 kb upstream and 1 kb downstream of the canonical TSS. Distal regions were defined as those within 1 Mb ± TSS, excluding promoter regions. Promoter peaks consisted of ATAC-Seq, H3K4me3, and H3K27ac peaks that reside in the promoter region. Distal peaks consisted of ATAC-Seq, H3K4me1, and H3K27ac peaks residing in distal regions, along with H3K4me3 peaks within 10 kb ± TSS. The following four link types were assessed: 1) distal peak to promoter peak, 2) distal peak to gene, 3) promoter peak to promoter peak, and 4) promoter peak to gene. In this analysis, 12,289 genes had at least one *promoter peak* and were included for further analysis. We conducted pairwise Spearman correlations between the variables identified in the described link types, leading to 12,481,099 unique tests. The *p*-values were adjusted using FDR (BH) within each link type and modality pair. We considered tests with an adjusted *p*-value < 0.05 to be statistically significant. 566,772 significant peak-peak or peak-gene relationships were identified where 10,786 genes are annotated via these significant correlations.

Due to variable relationships between distal peaks → promoter peaks and promoter peaks → genes, all peak → gene relationships were consolidated into “Pk2G links” with a consensus direction assigned using a prioritized, four-tier approach, where Tier 1 represents the highest confidence. Tier 1) Assign direction based on FDR significant links, bringing in correlation coefficients with nominal *p*-value < 0.05 to inform about direction where necessary. All peak:gene directions are in agreement. Tier 2) For links with relationships with conflicting direction and all links are FDR significant, the direction assigned to the link with the most significant FDR is honored. For example, if the distal peak → gene *r* > 0, the distal peak → promoter peak *r* > 0, and promoter peak → gene *r* < 0, the distal peak → gene relationship would return a positive relationship, while the distal peak → promoter peak →→ promoter peak → gene link would return a negative relationship. In this case, the distal peak → gene and distal peak → promoter peak FDRs would be compared. If the distal peak → gene FDR < distal peak → promoter peak FDR, then the consensus direction between the peak:gene would be positive. Tier 3) For links with conflicting direction, but not all links meet FDR significance, the relationship with the most significant nominal *p*-value < 0.05 was honored following a concept as in Tier 2. Tier 4) Direction of ambiguous peaks where the distal peak → promoter peak is FDR significant, but neither promoter peak → gene nor distal peak → gene correlations have a nominal *p*-value < 0.05, was assigned based on the correlation (*p*-value < 0.05) between the distal peak and other peaks significantly (FDR < 0.05) linked to the gene. The most FDR significant peaks were prioritized. Pk2G links with a direction confidence Tiers 1-3 were considered when identifying target genes in motif related analyses.

### SNP Calling from HCR/LCR RNA-Seq Data

Using the processed RNA-Seq BAM files, reads with an edit distance > 2 and duplicated reads were removed. Subsequently, BAM file base quality scores were recalibrated using the SplitNCigarReads, BaseRecalibrator, and ApplyBQSR commands from GATK^137^ version 4.3.0.0. Base recalibration used variants found in the eight founder strains of HCR/LCR as known sites^138^. SNPs were called using bcftools^139,140^ version 1.12 *mpileup* (-Q 20 -I -a DP,AD -d 10000 -E) and *call* (-mv -a GQ). HCR and LCR line assignments were provided as the population parameter for SNP calling. Genotypes with GQ < 35 were marked as missing and variants were removed if QUAL < 20 or > 50% of the genotypes were missing. The variant sites were further filtered to retain those discovered in the founder population^138^.

### Genotyping of F2 Population

A custom Affymetrix Axiom panel of ∼700,000 SNPs was designed to genotype the 613 F2 animals using DNA extracted from frozen liver tissue; SNPs in this panel were selected based on criteria previously described in Ren et al.^24^. These SNPs were based on the rn4 version of the rat genome assembly, then lifted over to the rn6 genome using LiftoverVcf (picard v2.8.1)^118^ and then filtered against an rn6 blacklist (https://github.com/shwetaramdas/maskfiles/blob/master/rataccessibleregionsmaskfiles/strains_intersect.bed), resulting in 380,990 autosomal SNPs. The genotypes were bidirectionally lifted over from the rn6 to the rn7 genome using GATK^137^ v4.3.0.0 *LiftoverVcf* (--RECOVER_SWAPPED_REF_ALT false). Variants were first assigned location-based IDs in the rn6 VCF using bcftools v1.7^140^ *annotate*(-x ID --set-id +’%CHROM\_%POS’), lifted to rn7, and then back to rn6. Variants that returned the same genomic position in the reverse (rn7 rn6) lift were retained using bcftools v1.7^140^ *view*. This resulted in 375,220 autosomal SNPs in the rn7 VCF. Genotypes were imputed from this panel for samples that have RNA-Seq data as well (described below).

### Genotype Imputation

#### Genetic Map Preprocessing

The rn7 genetic map published by De Jong et al.^141^ was used. Discrepancies throughout the file were addressed for autosomal chromosomes by the following preprocessing steps: 1) removal of sites where contig ≠ chr, 2) linear transformation of cM values for chromosomes with negative relationships between position and average cM, 3) removal of outlier cM:Position pairs, 4) sorted by position and smoothed the cM values by running mean weighted by standard deviation (bin size = 500). These steps removed 1,626 entries and resulted in a genetic map with 149,209 positions.

#### Imputation

IMPUTE5^142^ version 1.2.0 and SHAPEIT5^143^ version 5.1.1 were used to carry out genotype imputation. The reference panel published by Hermsen et al.^144^ was prephased using IMPUTE5 *xcftools_static view* (-m 0.03125). Target files were prephased using SHAPEIT5^143^ *phase_common_static*, providing the phased reference panel and referring to the preprocessed genetic map. Genomic regions were chunked for imputation using IMPUTE5 *imp5Chunker_v1.2.0_static*. Genotype imputation was carried out using IMPUTE5 *impute5_v1.2.0_static*, setting the effective population size to 52 and providing the phased reference panel, phased target file, preprocessed genetic map, and computed chunk and buffer regions. Post-imputation, genotypes were filtered to retain sites with a minor allele count (MAC) > 10 and posterior probability > 0.97 for all samples. As a quality control measure, imputed genotypes were compared to reference alleles derived from ATAC-Seq BAM files to verify consistency between the proportions of reference and alternate alleles at both homozygous and heterozygous sites (Fig. S16).

### Calculation of Fixation Index for HCR/LCR

The population differentiation between HCR and LCR lines was assessed by computing the fixation index (F_ST_)^145^ to compare reference allele frequencies (RAF). The following formula was used 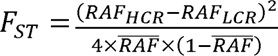. F_ST_ scores range from 0 to 1, where a score of 1 is interpreted as complete population differentiation. F_ST_ > 0.25 is described as very high differentiation^145^. A conservative threshold of F_ST_ > 0.3 is applied to identify 163,767 that exhibit high population differentiation between HCR and LCR.

### Comparison of F_ST_ to Peak Significance

We performed a linear regression for each modality using the model: 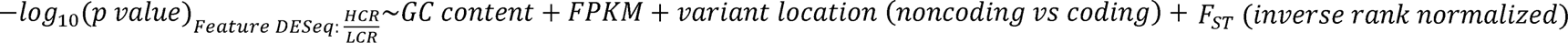. In instances where multiple variants reside within one epigenomic feature, the variant with the maximum F_ST_ value was retained to ensure unique epigenomic peak observations. GC content, FPKM, and variant location are included in the model to control for feature-specific artifacts.

### Motif Analysis

#### Motif Inclusion

The MotifDb^146^ R package was used to query motifs cataloged for humans and rats from CIS-BP, JASPAR, HOCOMOCO, and other databases. The initial 6,131 motifs were filtered to include TFs present in the RNA-Seq data. In instances of duplicate JASPAR motifs, the most recent version was kept. Redundancy was reduced by removing duplicated motifs (identical universal motif PPMs or TFBSTools::PWMSimilarity^147^ = 0), keeping only one instance of the particular motif per TF. This filtering resulted in 1,962 motifs from 516 transcription factors.

#### Preparation of Canonical Sequences

Sequences within significantly different peaks (defined in Fig. 2a) and ChromHMM regions (FDR < 0.1) were used as the input sequences for motif enrichment and PWM scans. Canonical HCR and LCR sequences were prepared for each feature set by replacing the reference allele with HCR and LCR alleles meeting the following criteria: 1) the variant site has an F_ST_ > 0.3 and 2) the alternate allele frequency within the given line is > 0.5. For regions where multiple SNPs fall within a close proximity (e.g., within the same feature), the one combination of ref/alt alleles that was most represented in the line was used for replacement. The rationale for this is to include relevant combinations of sites that indicate population differentiation while preventing over-representation of the input sequences in the analysis.

#### Haplotype-Aware PWM Scans

FIMO (v5.5.5)^148^ was used to carry out PWM scans for HCR and LCR sequences that extended ± 35 bp of a variant with F_ST_ > 0.3. Disrupted motifs are defined as those with Δ -log_10_(FIMO *p*-value) > 1.

#### Differential Motif Enrichment Analysis

Motif enrichment was performed, using AME (v5.5.5)^149^ with the arguments --control --shuffle scoring max, to specify the shuffled background model and scoring based on the maximum PWM match. The input sequences (pre-trimmed) were retrieved for each modality & direction of change (i.e., HCR up, LCR up) based on the following significance thresholds: A) ATAC-Seq peaks with FDR < 0.01 & abs(log_2_FC) > 0.5; B) H3K4me3, H3K4me1, and H3K27ac peaks with FDR < 0.1 & abs(log_2_FC) > 0.5; and C) ChromHMM states associated with open chromatin (EnhS_E1, EnhW_E3, EnhS_E4, PromWk_E5, PromA_E7, and PromPois_E8) with FDR < 0.1. For ChromHMM states, directionality was assigned by comparing mean posterior probabilities within significant regions between groups.

To control for differences in genomic territory associated with HCR up and LCR up regions within a given modality, we trimmed the larger set of features by removing the least significant regions until the length of sequences spanned within ∼1 kb of the smaller set of features. Final sequence lengths used for enrichment analysis per modality were: ATAC-Seq (872.3 kb), H3K4me3 (870.8 kb), H3K4me1 (2.5 Mb), H3K27ac (233.2 kb), EnhS_E1 (135.5 kb), EnhW_E3 (1.4 Mb), EnhS_E4 (1.3 Mb), PromWk_E5 (590.7 kb), PromA_E7 (339.5 kb), and PromPois_E8 (351.5 kb).

Motif enrichment was carried out independently for each modality and direction of change (e.g., HCR up and LCR up). For each modality and direction of change, two AME analyses were carried out using 1) HCR canonical sequences and 2) LCR canonical sequences.

Differentially enriched motifs were identified per modality using two approaches: a global comparison and an allele-specific comparison. In the global analysis, HCR sequences in HCR up regions were compared to LCR sequences in LCR up regions. For the allele-specific analysis, two comparisons took place: 1) HCR vs. LCR sequences in HCR up regions, and 2) HCR vs. LCR sequences in LCR up regions. For both global and allele-specific analyses, we compared the resulting AME -log10(*p*-values). Motifs were considered differentially enriched if the absolute difference in -log10(*p*-values) ranked in the top 10% of all Δ -log10(*p*-values) in the comparison.

In the allele-specific analysis, we identified regions that contributed toward the differential motif enrichment and contained motifs with disrupted PWMs. Starting with the AME results, regions were initially filtered to retain those that exclusively returned a true positive and either 1) were absent from the motif match in the counter group (e.g., when assessing HCR sequences, there was not a motif match returned in the LCR sequence), or 2) the difference in the PWM scores for the assessed region was greater than 1 (target group region – counter group region). The regions were further filtered to retain those that contained a variant with an F_ST_ > 0.3, following the assumption that regions without SNPs are likely to produce the same or similar PWM scores across different tests. Regions with disrupted motifs were further refined by directly comparing PWM scans of HCR and LCR sequences, retaining those with Δ -log_10_(FIMO *p*-value) > 1.

### Sample Swap Detection for F2 Analysis

QTLtools^150^ (v1.2) command mbv was used to detect sample swaps, using the F2 genotypes that were bidirectionally lifted over from rn6 to rn7. The flag --filter-mapping-quality 255 was applied when processing RNA-Seq BAM files, while this flag was not used for the pruned ATAC-Seq BAM files. When assessing ATAC-Seq sample swaps, matches with *n_het_covered > 5* were considered. Samples displaying a high degree of heterozygous matches were retained and sample ID swaps were corrected where necessary. One RNA-Seq and two ATAC-Seq samples were excluded due to low heterozygous consistency (<63%) with any genotype profile.

### ATAC-Seq Count Matrix Construction for F2 Analysis

To improve performance in the caQTL scan, a consensus peak matrix was uniquely prepared for the F2 samples. The peak count matrix was created by including peaks (narrow summits extended ± 150 bp) present in at least 20% of samples in either male or female groups. For overlapping peaks, the highest scoring peak was retained as the “candidate” region to create a non-redundant peak set. Inverse rank normalized counts, normalized for sex, were used for visualization.

### Sample Drop Criteria for F2 Analysis

We dropped outliers in PC space prior to performing the scans. We performed DESeq2^126^–style size factor normalization of the count matrices (as implemented in pyqtl, https://github.com/broadinstitute/pyqtl), dropped features where samples all had 0 counts, filtered to features with at least 10 normalized counts in at least 10 percent of samples, filled any remaining 0s with a value equal to (½ of the minimum non-zero value for that feature), log_10_ transformed the matrix, and then performed PCA on this normalized matrix. Outliers were determined based on visual inspection of the top 10 PCs. The final sample sizes for the (e/ca)QTL scans were 147 RNA-Seq samples and 132 ATAC-Seq samples.

### F2 eQTL & caQTL Analysis for Detection of eGenes and caPeaks

We performed cis-expression and chromatin accessibility quantitative trait locus scans (eQTL, caQTL) using the F2 RNA-Seq and ATAC-Seq data. We used a linear mixed model, implemented in GEMMA v0.98.5 (gemma -bfile $plink_prefix -n $phenotype_index -snps variants-to-test.txt -lmm -k $kinship_matrix -o $output_prefix -c covariates.txt)^151^. We computed the kinship matrix using GEMMA (gemma -bfile $plink_prefix -gk 1). For eQTL scans we tested all gene-variant pairs where the variant was within 1 Mb of the gene TSS. For caQTL scans we tested all peak - variant pairs where the variant was within 100 kb of the peak center. We normalized the count matrix as follows prior to the scan: 1) Filter to autosomal features, 2) Perform TMM normalization, 3) Filter to features with at least 5 (unnormalized) counts in at least 30% of samples, 4) Exclude features without any variants within the above-specified cis window, and 5) Inverse normal transform.

As covariates we included the top N gene expression / chromatin accessibility PCs (PCs calculated on the inverse normal transformed matrix), where N was selected in order to maximize the number of eGenes / caPeaks (this was determined by running (e/ca)QTL scans with 0, 5, 10, 15, … PCs, and then determining the number of eGenes / caPeaks yielded by each scan; the final N was 20 for RNA-Seq and 5 for ATAC-Seq).

After testing all feature - variant pairs, we performed two levels of multiple testing adjustment. First, we adjusted for the multiple tests performed against each feature using a slightly modified version of eigenMT^152^ (eigenMT was modified in order to collapse variants in perfect linkage disequilibrium before calculating eigenvalues; this modified version is available as git commit c89bf4e at https://github.com/porchard/eigenMT, and was run with arguments eigenMT.py --cis_dist $CIS_WINDOW --var_thresh 0.99 --window 10000 -- collapse-redundant-variants --CHROM $chrom --QTL qtl.txt --GEN $genotype_matrix --GENPOS $genpos -- PHEPOS $phepos --OUT $output). EigenMT estimates the number of independent tests per feature; we then multiplied the most significant observed *p*-value for each feature by the estimated number of tests (capping the product at 1, if greater than 1) to get a feature-level *p*-value. We then adjusted for the multiple testing across features, performing a Benjamini-Hochberg correction^130^ on the feature-level *p*-values to get an FDR for each feature. We then applied an FDR threshold of 5% to determine whether each gene or peak had a significant (e/ca)QTL. For each eGene / caPeak, the variant with the most significant *p*-value was set as the eVariant / caVariant (due to linkage disequilibrium, in many cases multiple variants had the same *p*-value; for downstream analyses sensitive to this, we randomly selected one of these variants to be the representative eVariant / caVariant). Alternative handling of linkage disequilibrium from the significant results included grouping (e/ca)Variants with r^2^ ≥ 0.99 to the most significant variant, labeled as (e/ca)VariantClusters.

### Testing Concordance Between F2 Allelic Effects and HCR/LCR Differences

To test whether the allelic effects detected in the segregating F2 population account for the epigenomic and transcriptomic differences observed between the HCR and LCR lines, we compared the direction of effect for each measurement across the two studies. Given that the two studies are statistically independent and are from separate cohorts, agreement in the direction of effect is evidence that the line differences are driven by heritable regulatory variation rather than by line-specific technical or environmental effects.

Analyses were restricted to variants with F_ST_ > 0.3 between HCR and LCR and three comparisons were made with unique pairing rules:

1. *F2 caQTL effects compared to HCR/LCR differential chromatin accessibility and histone modification abundance*. HCR/LCR peaks from modalities associated with open chromatin (ATAC-Seq, H3K27ac, H3K4me1, and H3K4me3) were overlapped with F2 caPeaks. Variant-peak interactions were refined to caPeaks where the associated variant was within the peak or in strong linkage disequilibrium (r^2^ > 0.5) with the most significant caVariant for that peak. HCR/LCR differential results were signed by DESeq log_2_FoldChange. In the F2 caQTL analysis, the effect allele was defined as the allele with the greater frequency in HCR and resulting -log_10_(*p*-values) were signed by slope. Independent observations were obtained by grouping variants into their caVariantClusters for testing and visualization.
2. *F2 eQTL effects were compared with HCR/LCR differential gene expression*. Genes with HCR/LCR DESeq FDR < 0.01 and eGenes where a variant was in strong linkage disequilibrium (r^2^ > 0.9) with the top eVariant were retained. HCR/LCR differential results were signed by DESeq log_2_FoldChange. In the F2 eQTL analysis, the effect allele was defined as the allele with the greater frequency in HCR and resulting -log_10_(*p*-values) were signed by slope. The overlapping genes were distinct and treated as the unit of observation.
3. *F2 eQTL effects were compared with the significance of HCR/LCR Pk2G links*. Significant Pk2G regions were overlapped with variants and joined to the eQTL results by variant (r^2^ > 0.9 with the lead eVariant) and target gene. Pk2G links were signed by the direction of relationship. For the F2 eQTL results, the effect allele was coded per variant-peak pair to the allele that increased the paired peak signal (e.g., HCR allele for HCR up peaks, and the LCR allele for the LCR up peaks). This aligns the eQTL effect with the allele driving the regulatory element, allowing the direction of the variant’s effect on its target gene to be compared with the direction of the Pk2G link. Variants were grouped into their eVariantClusters to obtain independent observations for testing and visualization.

Following these comparisons, points in quadrants I and III in Fig. 5c-e agree between the two studies. Directional concordance was tested using a one-sided binomial test (expected = 50%, alternative = “greater”).

## Supporting information

Supplemental Protocol S1

## Abbreviations

AME: analysis of motif enrichment
BCAA: branched-chain amino acid
caQTL: chromatin accessibility quantitative trait locus
CRF: cardiorespiratory fitness
ETF: electron-transfer flavoprotein
ETS: E26 transformation-specific
eQTL: expression quantitative trait locus
FDR: false discovery rate
F_ST_: fixation index
GSEA: gene set enrichment analysis
HCR: high capacity runners
LCR: low capacity runners
PCA: principal component analysis
Pk2G: peak-to-gene
PWM: position weight matrix
RPM: reads per million
SNP: single nucleotide polymorphism
TCA: tricarboxylic acid
TF: transcription factor
TSS: transcription start site
VO_2_max: maximal oxygen uptake
VST: variance-stabilizing transformation

## Acknowledgements

This research was supported in part through computational resources and services provided by Advanced Research Computing (ARC), a division of Information and Technology Services (ITS) at the University of Michigan, Ann Arbor. Next-generation sequencing was carried out in the Advanced Genomics Core at the University of Michigan. Dr. Nathan Xi and the University of Michigan Animal Metabolic, Physiological, & Behavioral Phenotyping Core assisted in the animal studies. Johanna Y. Fleischman, Gayatri R. Iyer, Martin J. Walsh, and Alexander J. Weitzel for reviewing the manuscript and providing advice as it was being prepared.

## Funding

Funding for the collection of HCR/LCR skeletal muscle samples was provided by a pilot award to CRE from the Michigan Nutrition Obesity Research Center, supported by National Institutes of Health grant P30 DK089503

A. Alfred Taubman Medical Research Institute (CFB)

National Institutes of Health grant U24 DK112342 (CFB)

National Institutes of Health grant R01 DK117960 (SCJP)

University of Michigan Systems and Integrative Biology Training Grant T32 GM008322 (AMW)

The LCR-HCR rat model was funded by National Institutes of Health Office of Research Infrastructure Programs grant P40OD-021331 (LGK, SLB)

## Author contributions

Conceptualization: AMW, CRE, JZL, SCJP, CFB

Methodology: AMW, PO, CRE, SLB, LGK, SCJP, CFB

Validation: AMW, PO, SCJP, CFB

Formal analysis: AMW, PO

Investigation: AMW, CRE, NM, MKT

Resources: SLB, LGK, JZL, CFB

Data curation: AMW, PO

Writing – original draft: AMW, SCJP, CFB

Writing – review & editing: all authors Visualization: AMW

Supervision: PO, JZL, SCJP, CFB

Project Administration: MKT, CRE, SCJP, CFB

Funding Acquisition: JZL, SCJP, CFB

### Competing interests

The authors declare no competing interests.

### Data, code, and materials availability

This study did not generate new unique reagents.

Raw (FASTQ files) and aligned (BAM files) sequencing data will be deposited on NCBI GEO upon final submission. The HCR/LCR posterior probability matrices, HCR/LCR and F2 count matrices, HCR/LCR and F2 imputed genotypes, and the F2 Affymetrix Axiom panel will be publicly available by uploading to appropriate database by the date of publication. Processed data and results referenced in this manuscript are available in a Zenodo repository (DOI https://doi.org/10.5281/zenodo.19699175).

All original code will be deposited at a Zenodo repository to be publicly available by the date of publication.

Any additional information required to reanalyze the data reported in this paper is available from the corresponding author upon request.

## Supplemental Figures

**Fig. S1:**
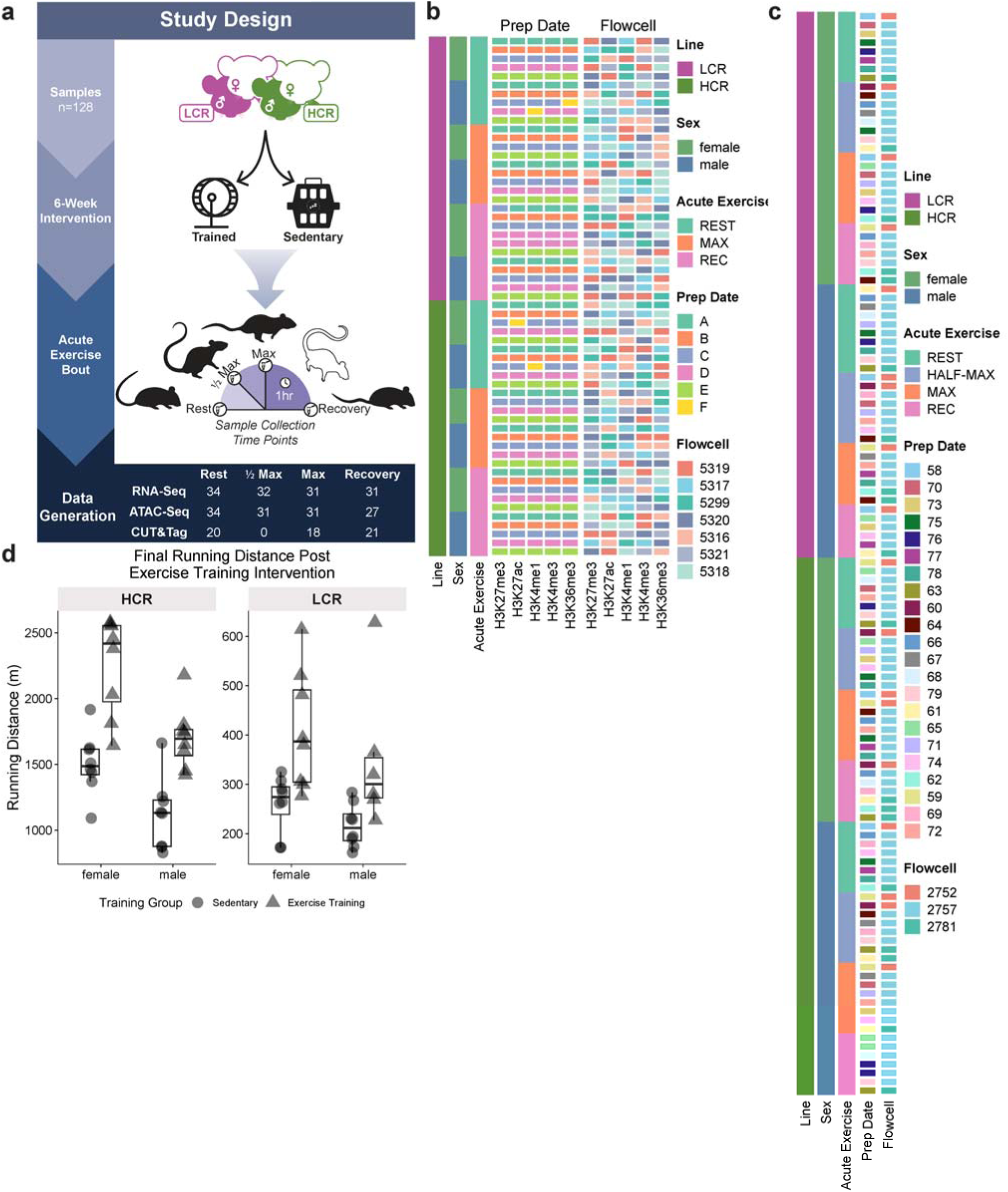
Detailed Breakdown of Study and Randomization, related to Fig. 1. (**a**) Study design that included 128 HCR and LCR rats, balanced for sex. Half of the cohort underwent a six-week endurance training regimen while the remaining rats were sedentary for that same time period. Rats were split into four sample collection time groups and subjected to an acute exercise bout with subsequent tissue collection either prior to, during, or after a near-exhaustion acute exercise bout. High throughput sequencing libraries were generated from gastrocnemius muscle, with the number of libraries produced per acute exercise bout group provided. Note that CUT&Tag libraries were not produced for samples in the half-max group, and fewer CUT&Tag libraries were produced for the remaining acute exercise bout groups. (**b**) Randomization schema of CUT&Tag libraries. (**c**) Randomization schema of ATAC-Seq libraries. (**d**) To assess the effect of the exercise training, we performed a linear regression using the formula *final running distance ∼ *line* + *sex* + exercise training group*. The results showed that running distance significantly increased following training (β = 404 m, *p*-value = 2.97×10^-07^).

**Fig. S2:**
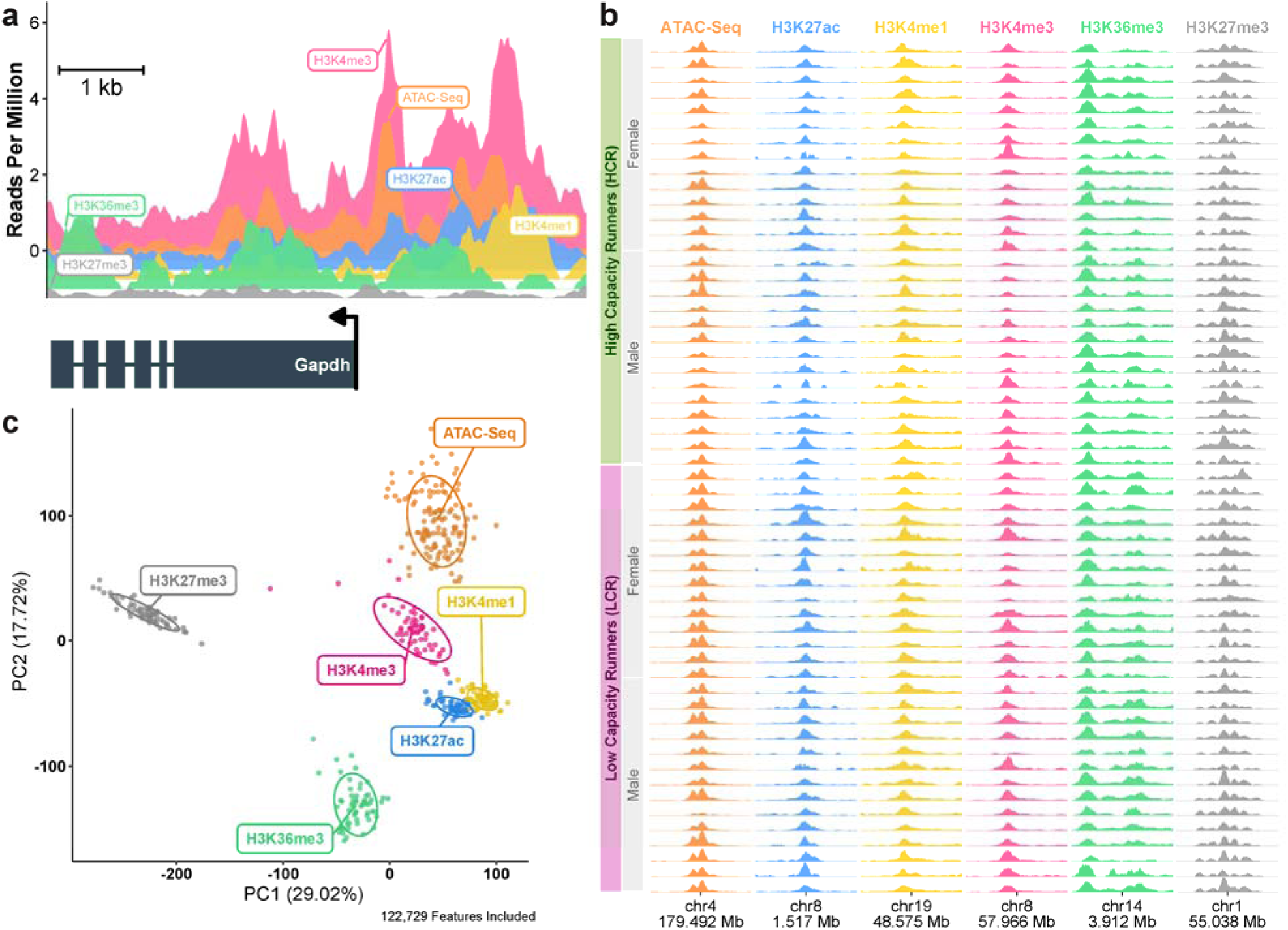
Data Overview, related to Fig. 1. (**a**) Example signal tracks at the *Gapdh* locus in a single sample (rat R18804), showing from back to front, H3K4me3, ATAC-Seq, H3K27ac, H3K4me1, H3K36me3, and H3K27me3. *Gapdh* is transcribed on the negative strand. (**b**) Signal reproducibility across the 56 samples with full epigenomic data (refer to Fig. S1a for details pertaining to library generation). Each column shows one modality at a different representative locus, displayed as the maximum RPM within 10 bp bins, smoothed using a rolling mean (bin size = 50). Each column represents a 4 kb genomic window centered around the midpoint annotated on the x-axis. Y-axis ranges are fixed within each column with a minimum value of zero and modality-specific maxima: ATAC-Seq (9.9 RPM), H3K27ac (5 RPM), H3K4me1 (10.3 RPM), H3K4me3 (15.4 RPM), H3K36me3 (6.6 RPM), and H3K27me3 (6.1 RPM). The presence and abundance of signal across genomic bins are consistent within modality. (**c**) PCA of genome-wide signal within 10 kb bins (inverse rank normalized, blacklist filtered). Sequencing libraries cluster by modality, demonstrating reproducibility across samples.

**Fig. S3:**
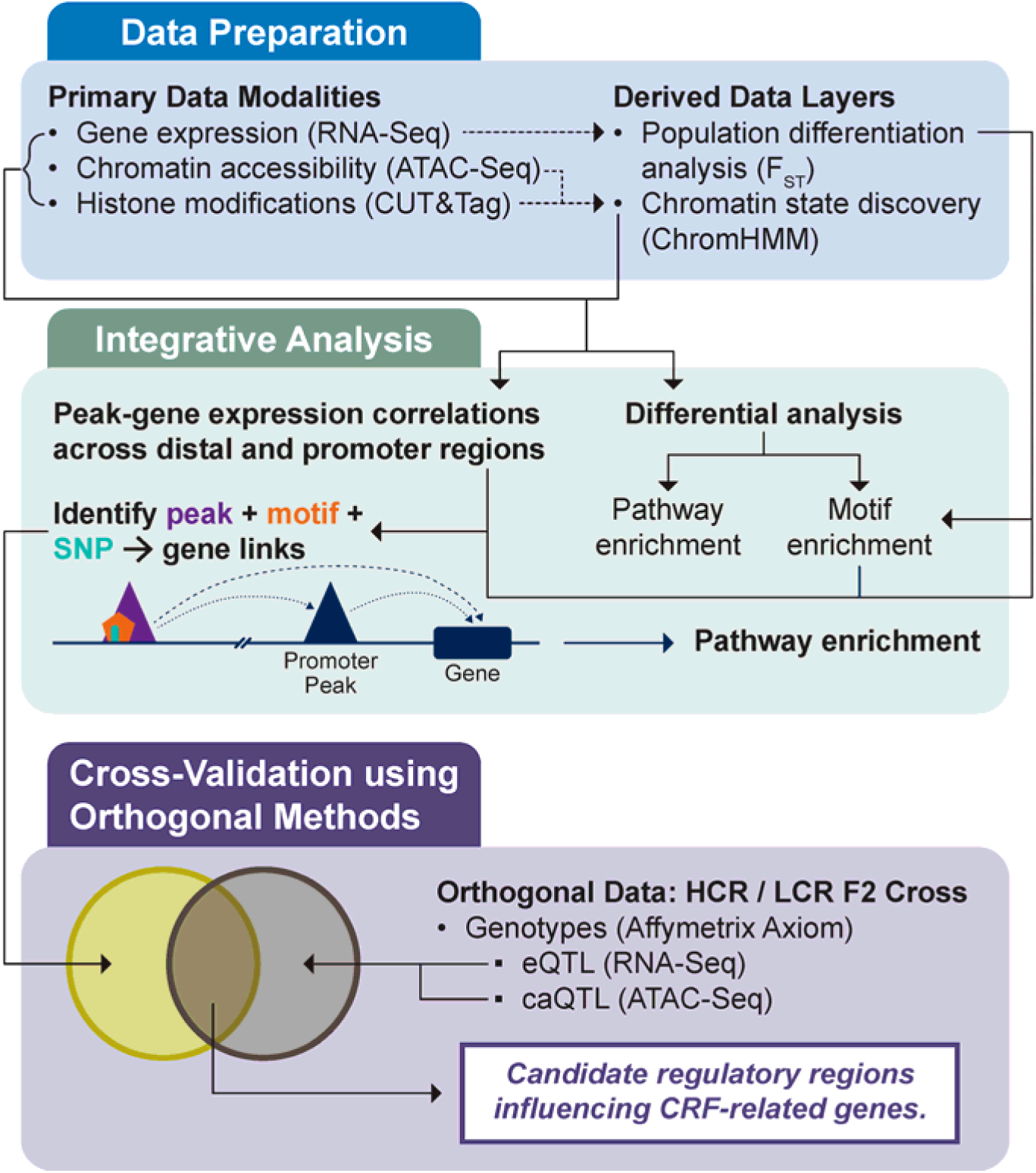
Multi-Modal Data Framework, related to Fig. 1. Overview of the data modalities included in the study, categorized by whether the data used is primary or derived. Primary data modalities are sequencing libraries prepared from skeletal muscle tissue. Derived data layers were generated by utilizing the primary data modalities. The green box shows the workflow in which datasets are analyzed independently and then integrated to identify variant:peak:gene links. The purple box presents the concept of the validation study.

**Fig. S4:**
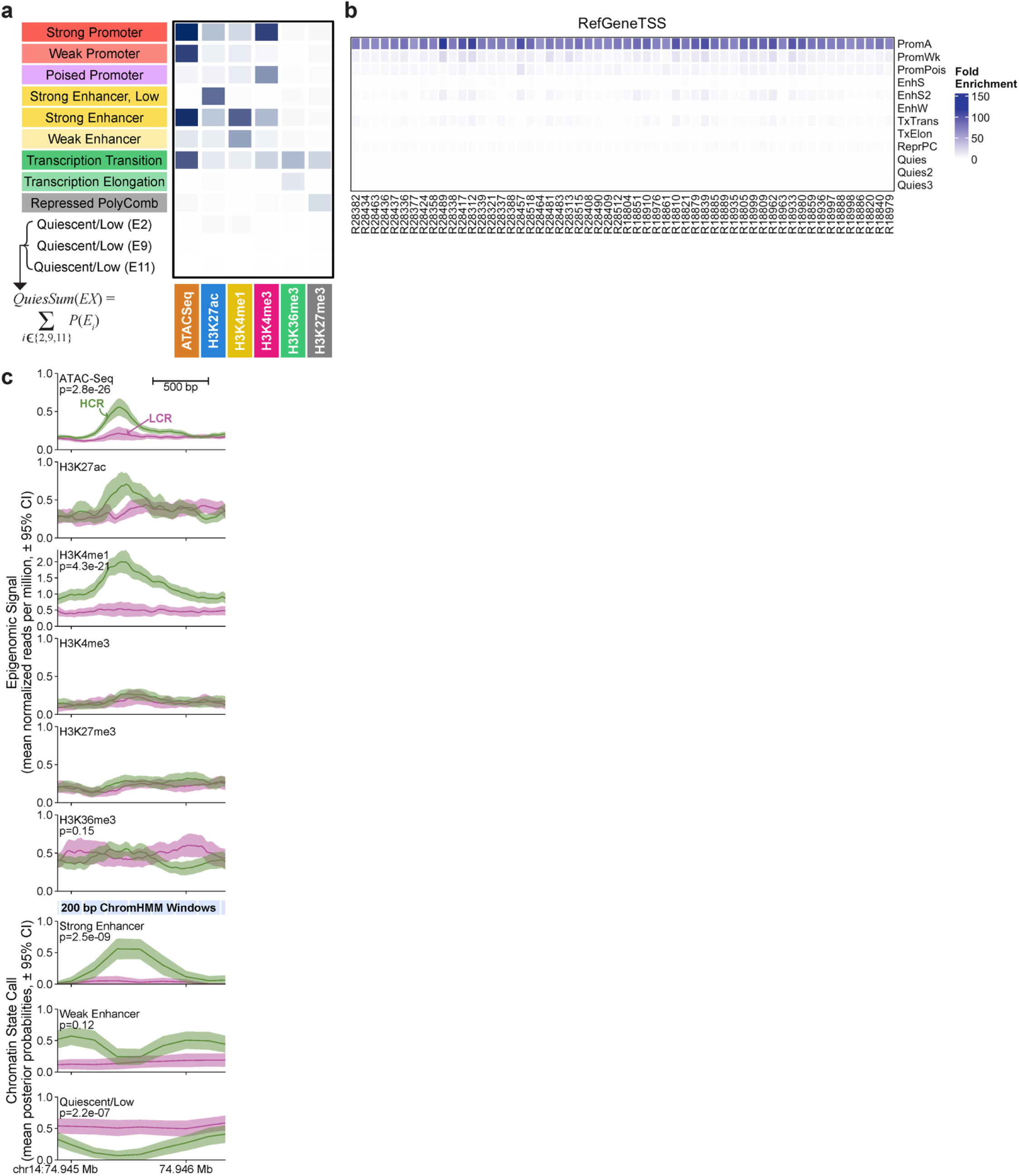
ChromHMM Analysis, related to Fig. 2. (**a**) ChromHMM emission plot showing chromatin state definitions where darker blue indicates a greater probability of mark presence in the emission state. Three low-signal states were summed into a single ‘Quiescent/Low’ state. (**b**) QC plot from ChromHMM analysis, showing the enrichment for the Strong Promoter chromatin state at TSSs across the genome. The likelihood of ATAC-Seq and H3K4me3 presence associated with the Strong Promoter state is expected at TSSs. (**c**) Example locus on chr14 showing normalized ATAC-Seq and CUT&Tag counts (reads per million) and ChromHMM posterior probabilities across 59 HCR (green) and LCR (magenta) rats. For each sample, ATAC-Seq and CUT&Tag signal was smoothed using a rolling mean applied to the maximum signal within 10 bp bins (bin size = 5), and values were summarized by plotting the mean ± 95% CI across samples. Posterior probabilities (mean ± 95% CI) are plotted at the midpoint of each 200 bp ChromHMM window, which represent the likelihood of the chromatin state occurring at these loci. Increased ATAC-Seq and H3K4me1 signal in HCR coincides with a rise in Strong Enhancer and dip in Quiescent/Low state probability, indicating a clear chromatin state difference between HCR and LCR. *P*-values from differential analyses are printed on each panel (not shown if not tested). Notably, ATAC-Seq libraries were prepared independently from CUT&Tag libraries and show highly concordant signals across ATAC-Seq and histone marks associated with open and active chromatin, further supporting the reproducibility and validity of these data.

**Fig. S5:**
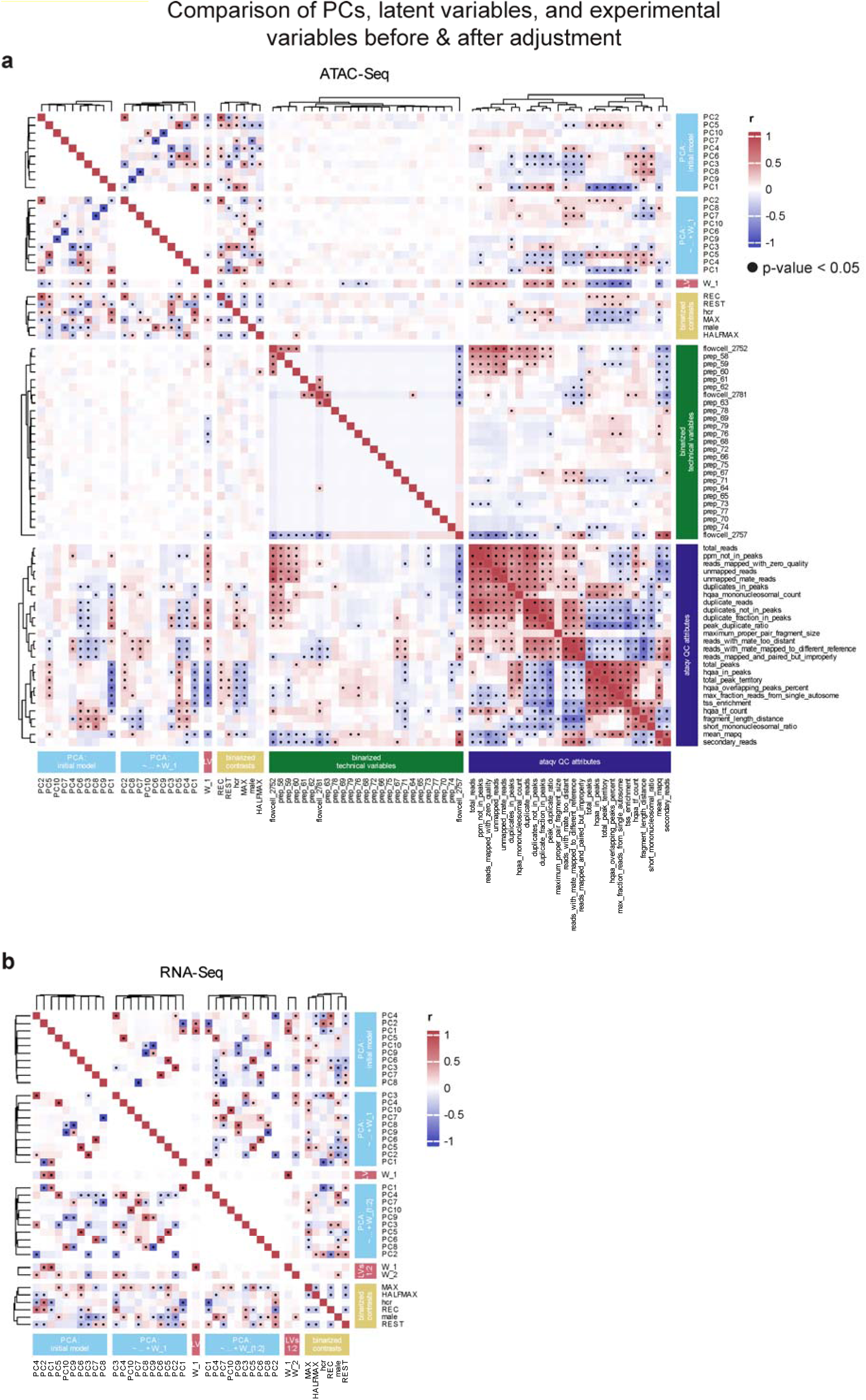
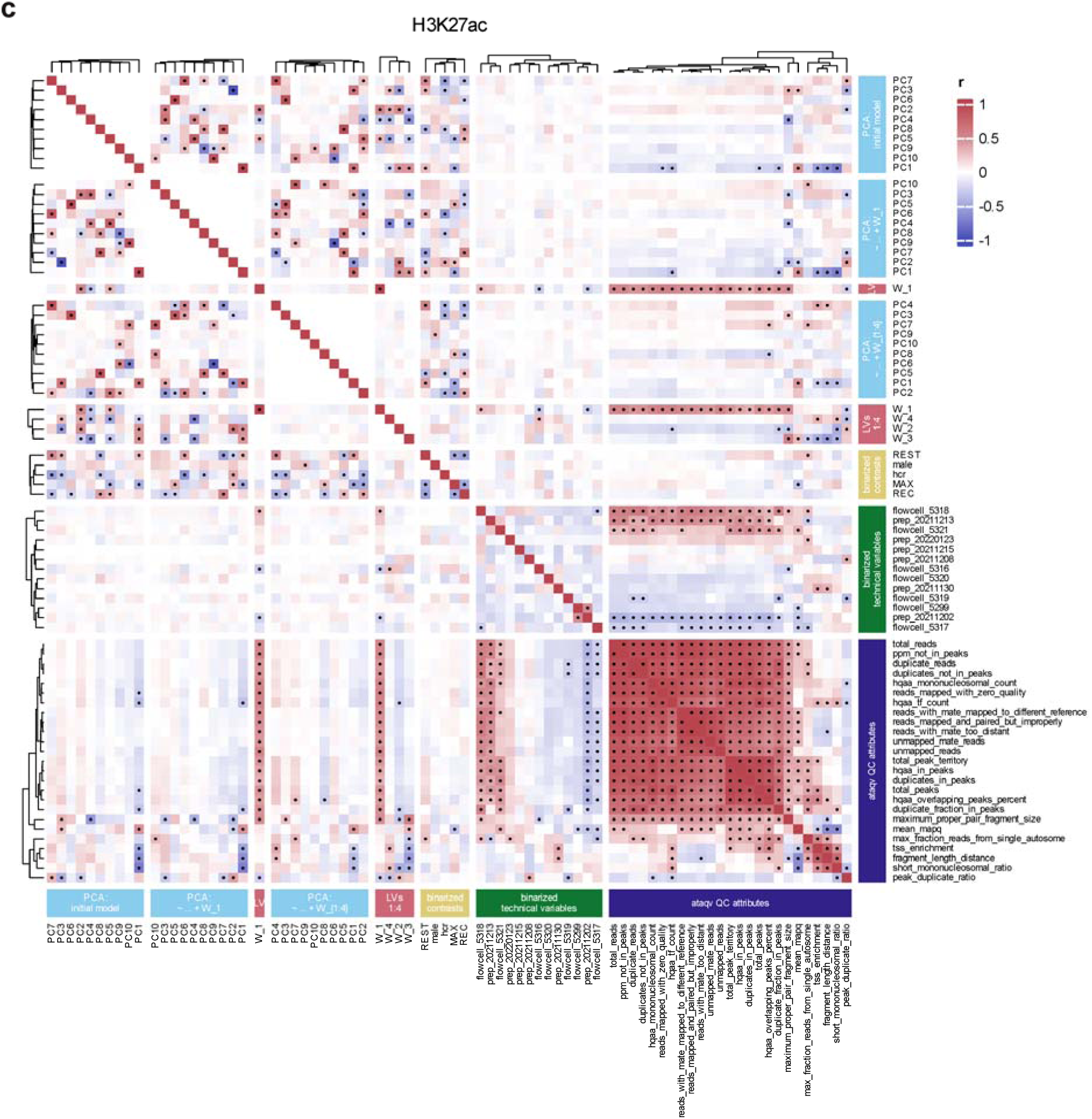
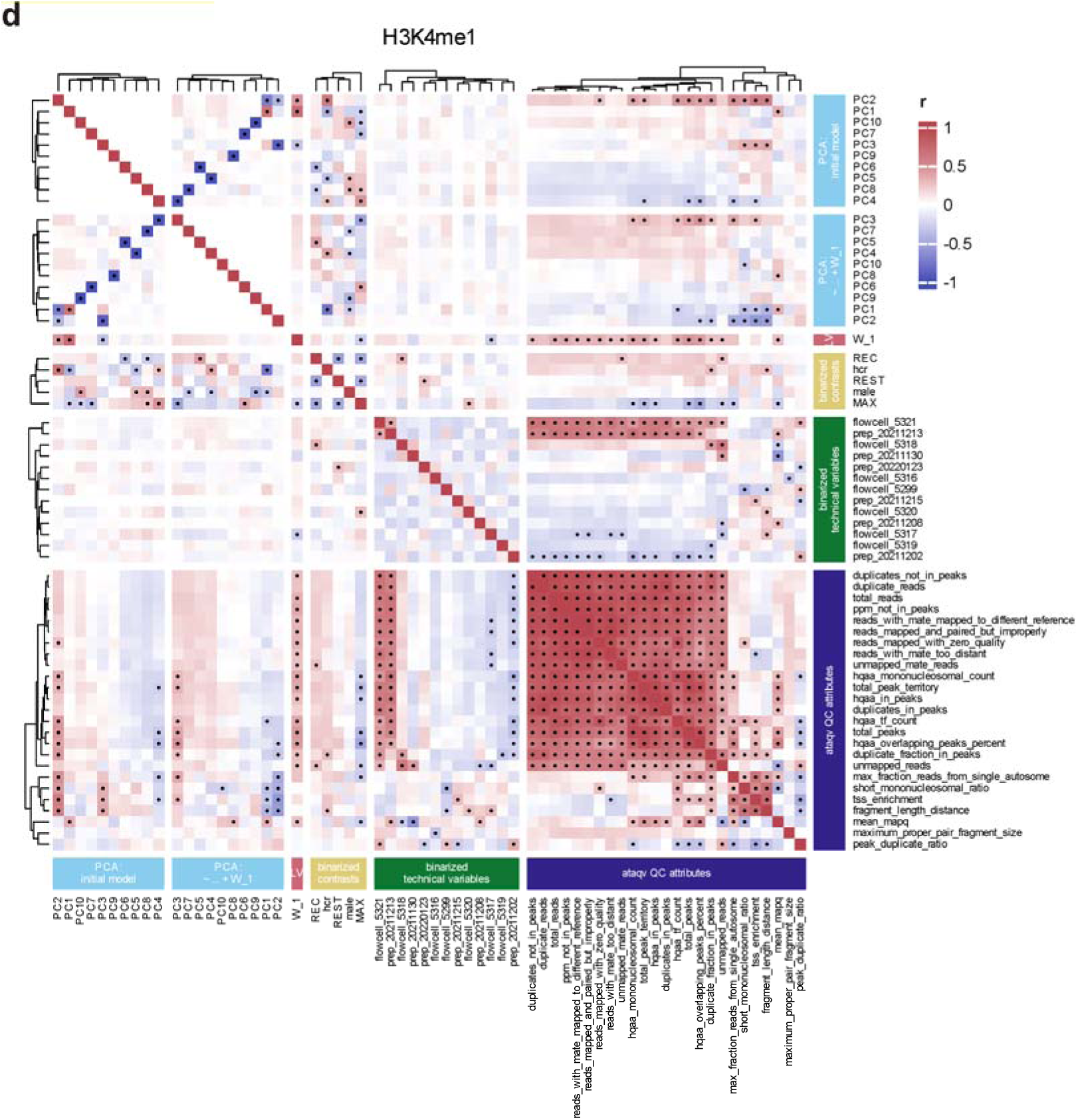
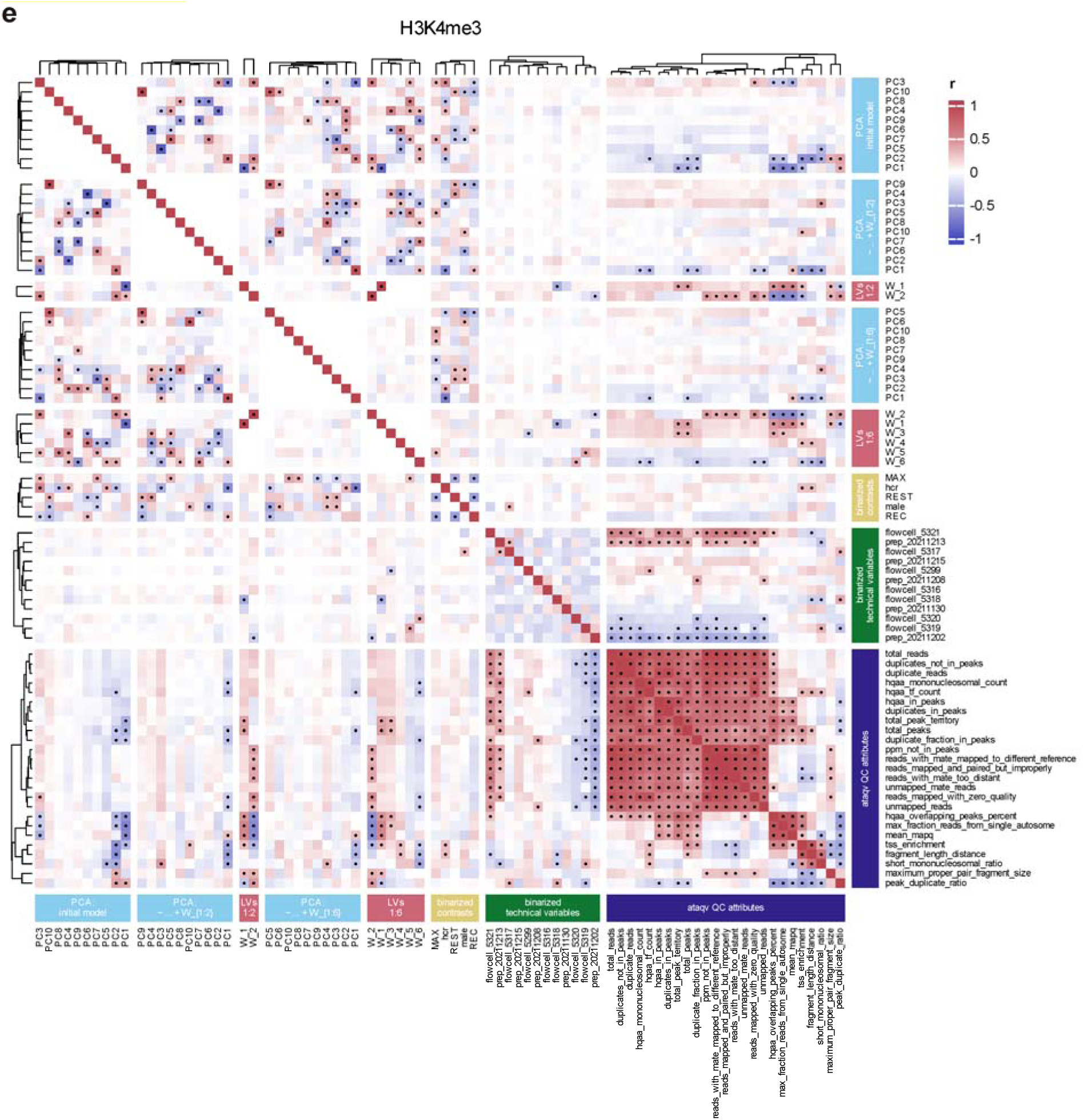
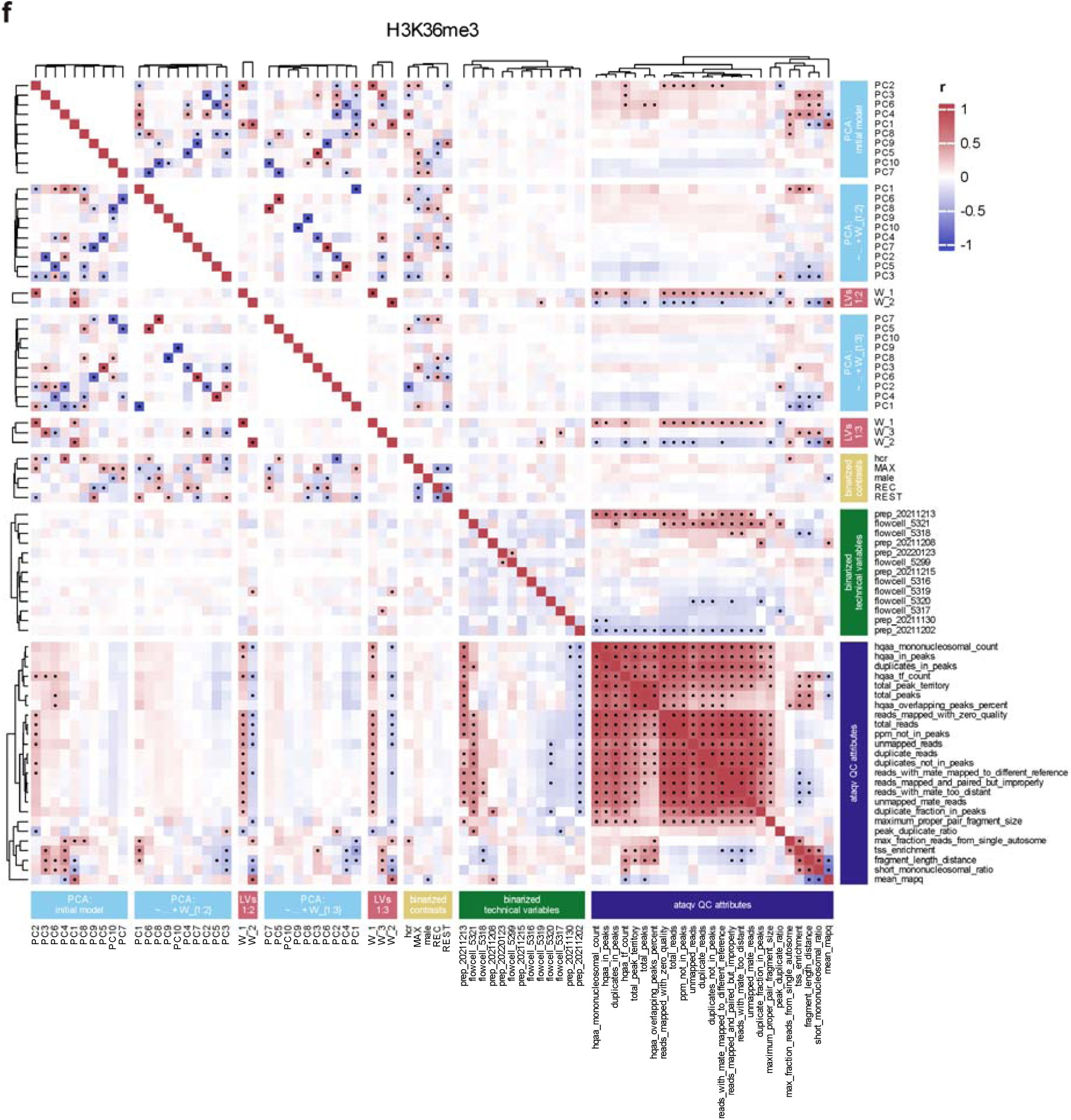
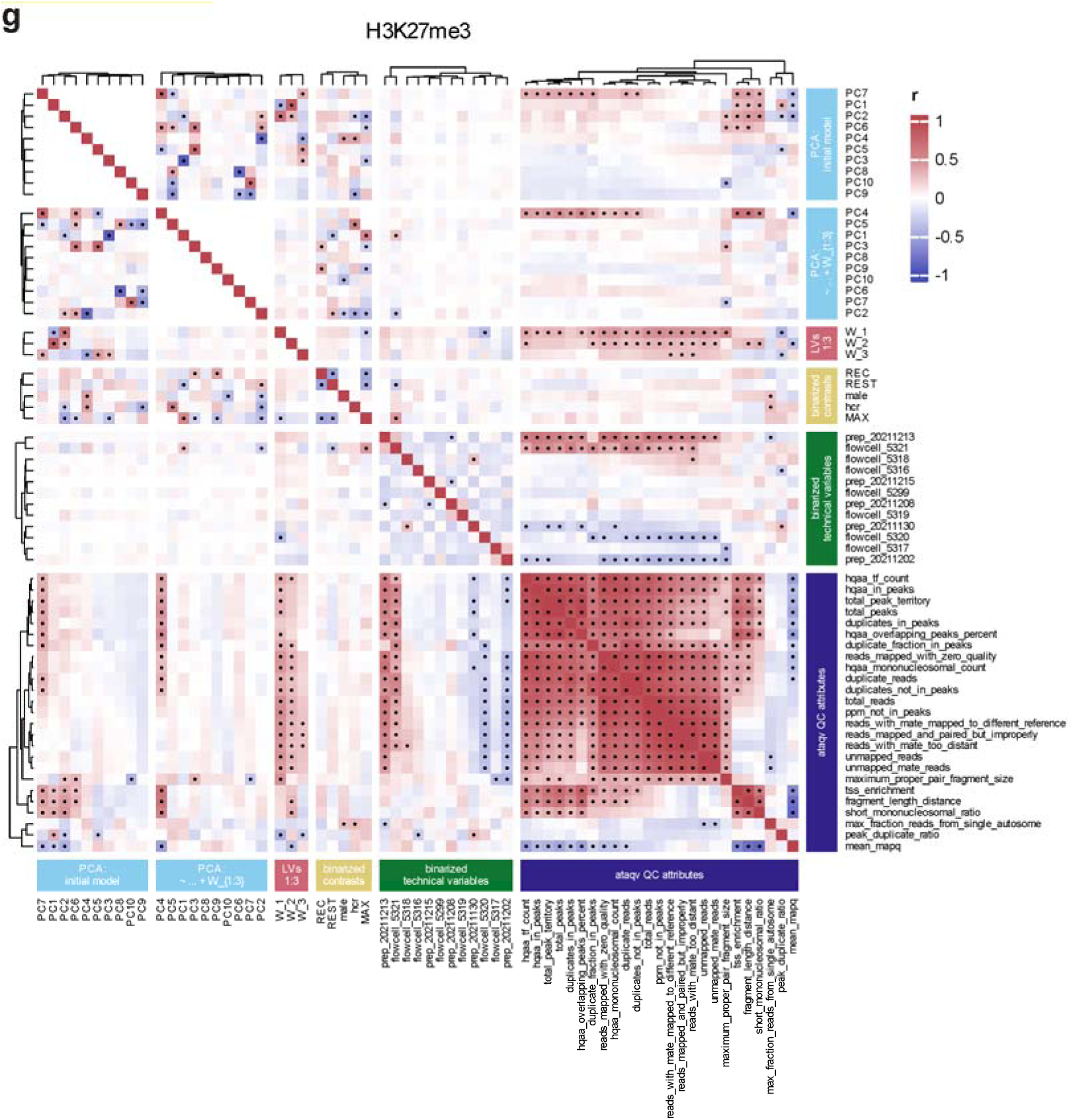
Comparison of PCs, Latent Variables, and Experimental Variables Before and After Adjustment of ATAC-Seq, CUT&Tag, and RNA-Seq Data, related to Fig. 2. (**a-g**) Heatmap of correlation between binarized experimental contrasts (yellow), considered latent variables (red), and PCs before and after inclusion of latent variables (light blue), with each other and QC metrics associated with epigenomic sequencing data (dark blue).

**Fig. S6:**
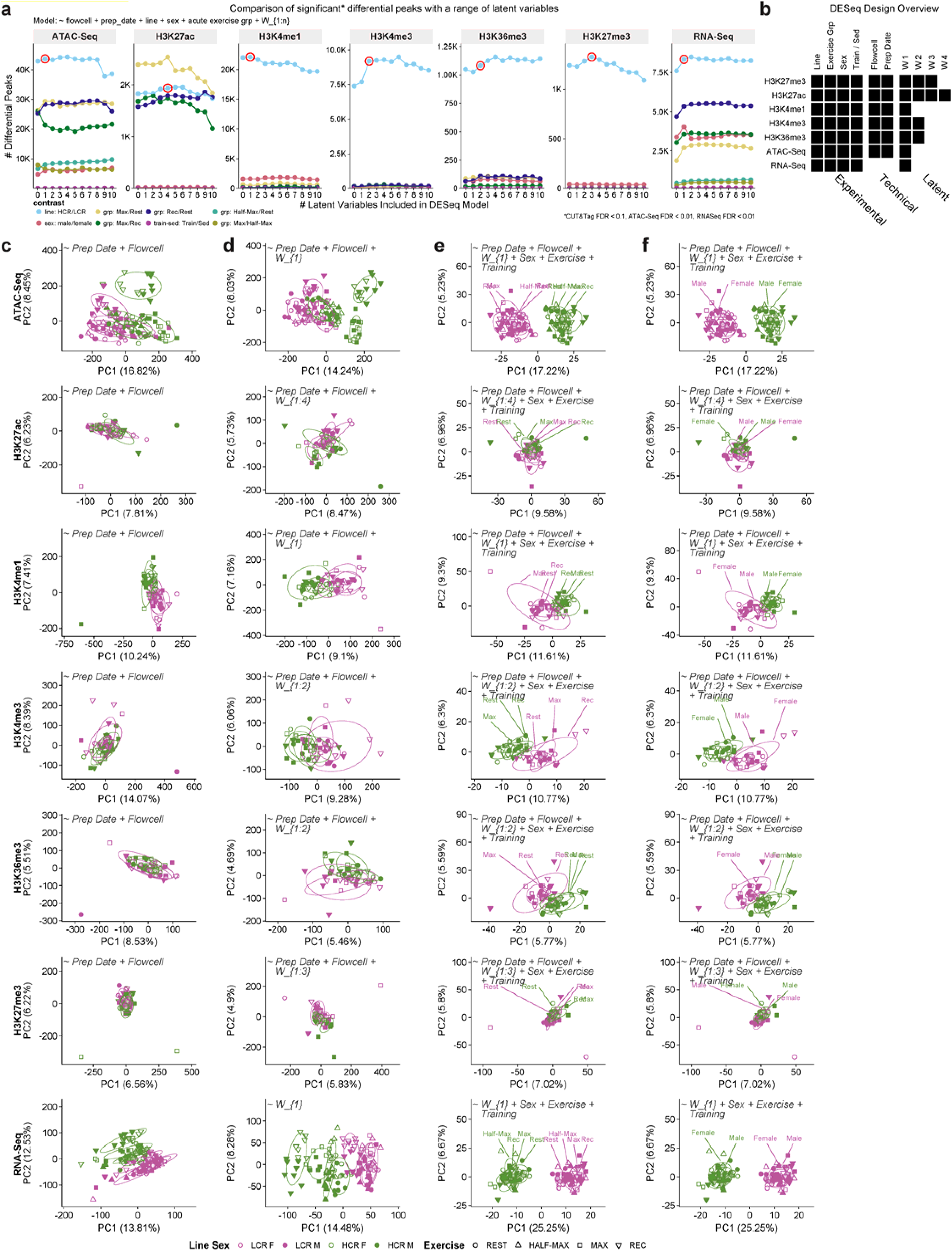
Identification of Latent Variables, DESeq2 Models, and Variation of Data Before and After Adjustment, related to Fig. 2. (**a**) Comparison of the count of significant differential peaks (y-axis) resulting between all experimental contrasts (colored) when a range of latent variables are included in the DESeq model (x-axis). The number of latent variables that we chose to include in all subsequent models (i.e., DESeq, adjusting of VST transformed counts for correlation and data visualization) for each modality is circled in red. (**b**) DESeq design overview for each modality subject to differential analysis by this method. (**c**) PCA plots of VST transformed counts for each modality normalized for technical variables by adjusting based on the formula printed on each plot. (**d**) PCA plots of VST transformed counts for each modality normalized for technical and latent variables (formula printed on the plot). This shows how outliers move towards their groups, improving group clustering by normalizing for unknown variation. Additionally, these plots show how strong the line effect is relative to acute exercise and sex prior to the removal of this effect. (**e-f**) VST transformed counts where the line effect was isolated by adjusting for additional experimental (sex, acute exercise, and training) and technical variables (prep batch, sequencing batch, latent variables) via linear regression. The linear model included the line variable to protect this effect. These plots confirm that acute exercise (**e**) and sex groups (**f**) are centered post-removal of the additional experimental and technical contrasts, exemplifying successful isolation of the HCR/LCR effect in this data. Centering of training samples is not displayed because that experimental contrast resulted in minimal molecular changes.

**Fig. S7:**
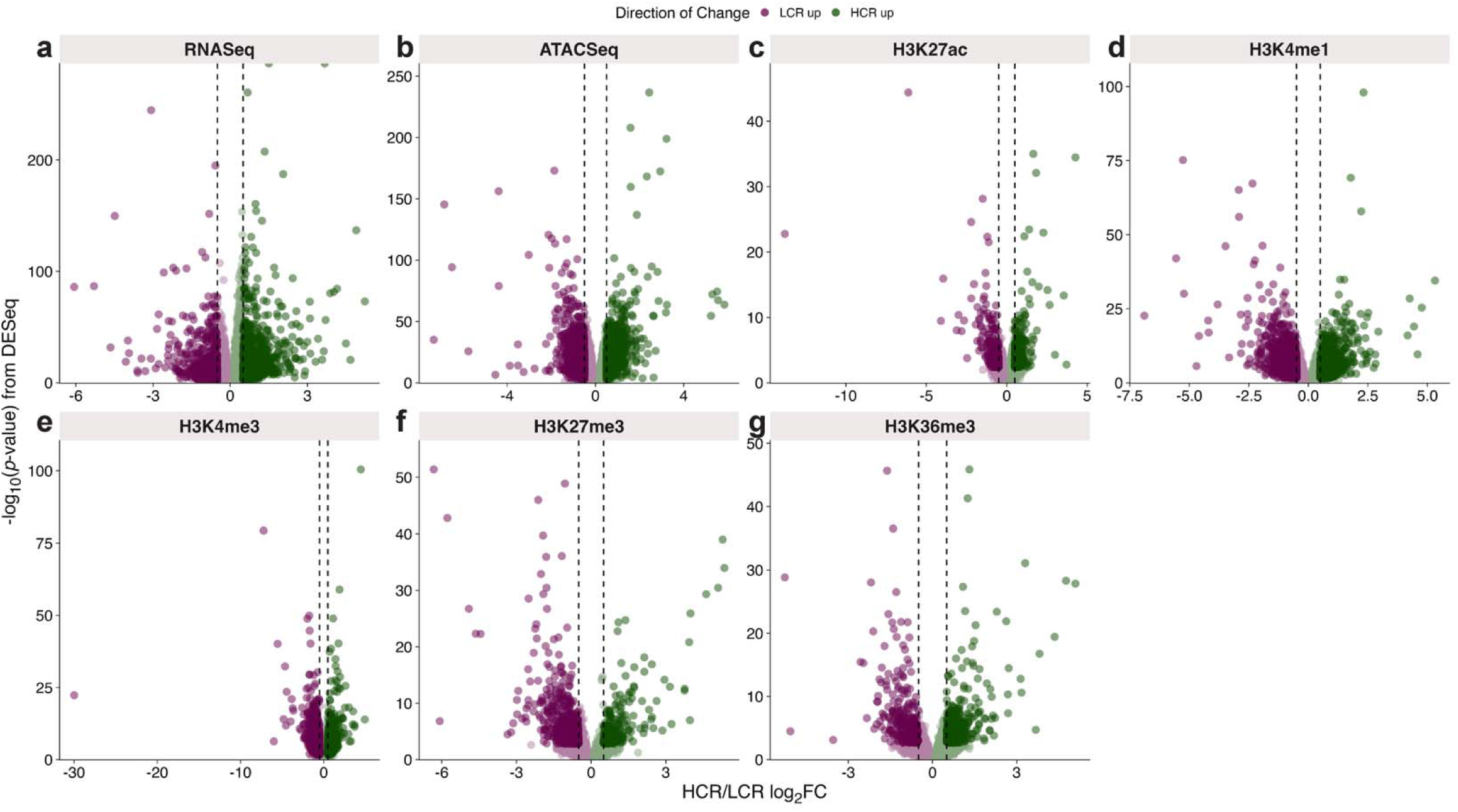
Differential Chromatin Accessibility, Histone Modification Abundance, and Gene Expression, related to Fig. 2. Volcano plots created from the HCR/LCR DESeq output, with emphasis on the regions that pass the significance threshold provided in Fig. 2a for each sequencing-based modality: (**a**) RNA-Seq (gene expression), (**b**) ATAC-Seq, (**c**) H3K27ac, (**d**) H3K4me1, (**e**) H3K4me3, (**f**) H3K27me3, (**g**) H3K36me3.

**Fig. S8:**
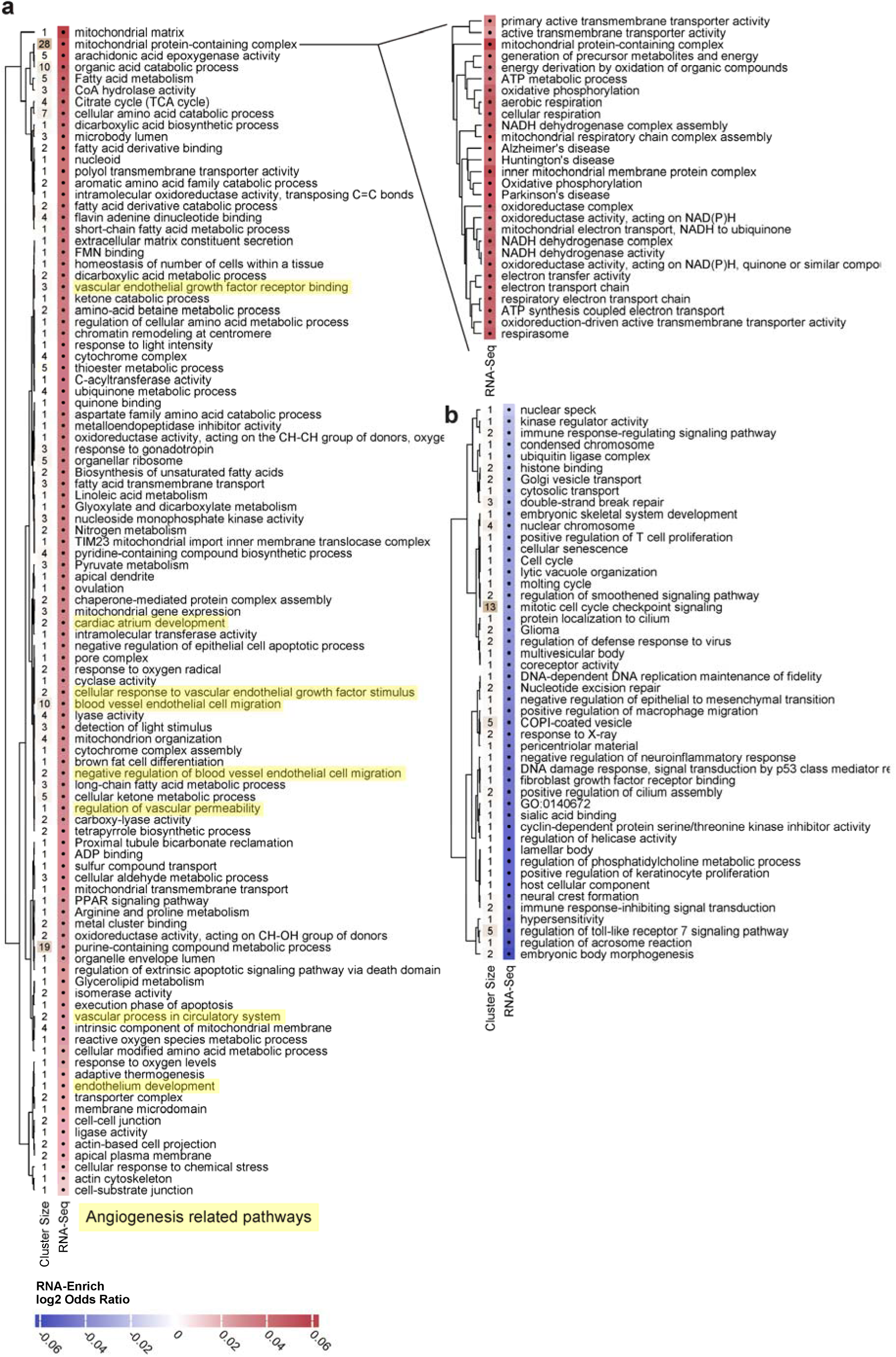
Differential RNA-Seq Pathway Enrichment Results, related to Fig. 2. RNA-Seq pathway enrichment analysis. HCR/LCR differential gene expression significantly enriched for 342 gene sets via RNA-Enrich analysis (FDR < 0.05), which were collapsed into 149 representative clusters and plotted separately by upregulation in HCR (**a**) and LCR (**b**). The rows are hierarchically clustered based on the log_2_ odds ratio of the RNA-Enrich results plotted. The number of gene sets within the hierarchical cluster is annotated on the left-hand side of the heatmap. The largest cluster in panel (**a**), mitochondrial protein-containing complex, is expanded upon to the right to show the individual terms that are part of that cluster. Angiogenesis-related clusters are highlighted (FDRs = 6.9×10^-5^ to 3.6×10^-2^).

**Fig. S9:**
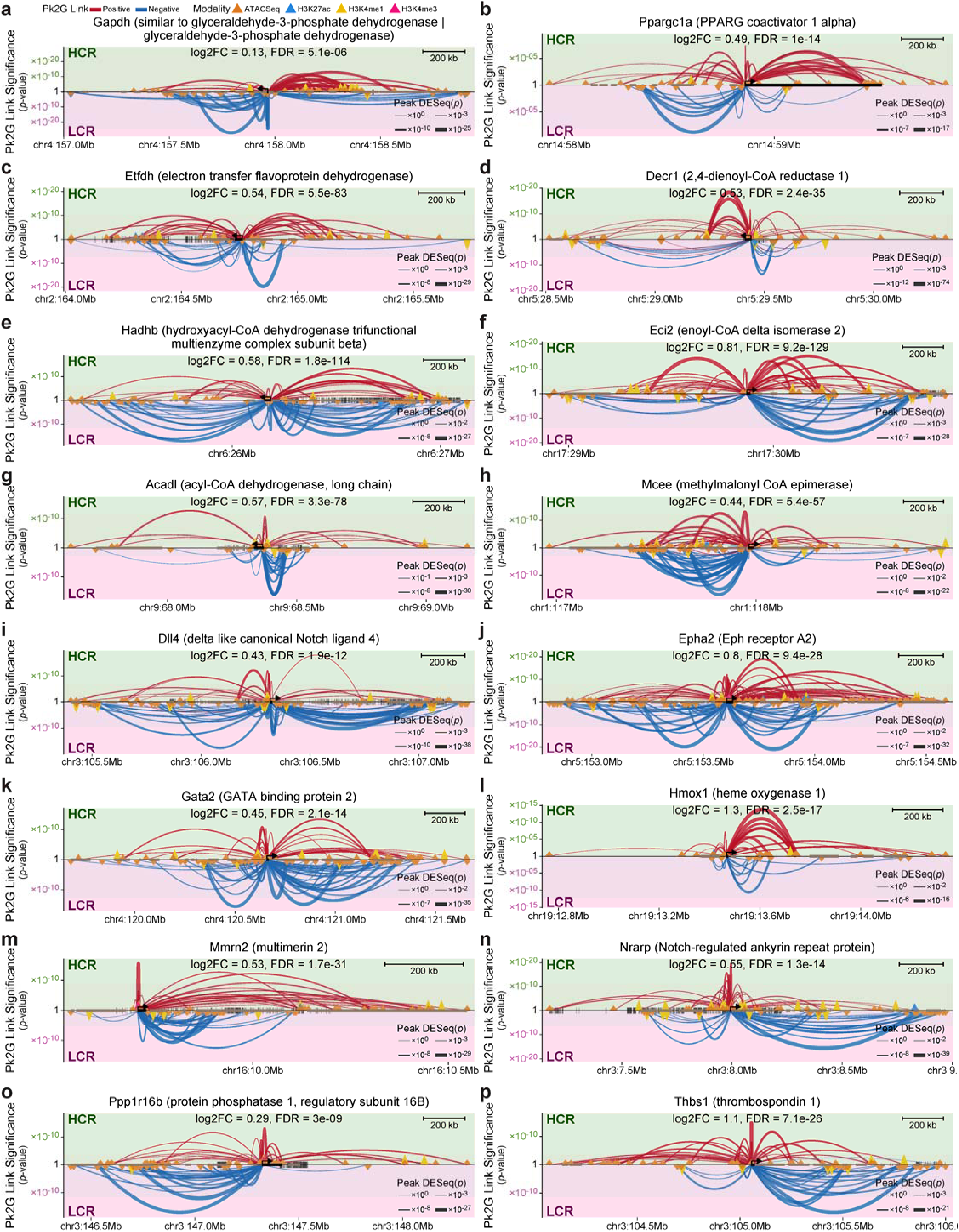
Pk2G Examples, related to Fig. 3. Peak-gene regulatory architecture surrounding (**a**) housekeeping gene *Gapdh*, (**b**) mitochondrial biogenesis regulator *Ppargc1a*, (**c-h**) fatty acid and BCAA oxidation genes, and (**i-p**) angiogenesis-related genes. Regulatory elements (triangles, colored by modality) are positioned by genomic distance from the TSS (arrow) of the gene (black rectangle). Peaks with DESeq log_2_FC > 0 (HCR up) are located above the midline, and LCR up peaks are shown below. Arc height denotes the significance of the Pk2G links, colored by direction of Pk2G link, with linewidth proportional to DESeq significance of the regulatory element. Vertical ticks at the midline represent SNPs with F_ST_ > 0.3 and additional rectangles represent other genes within view.

**Fig. S10:**
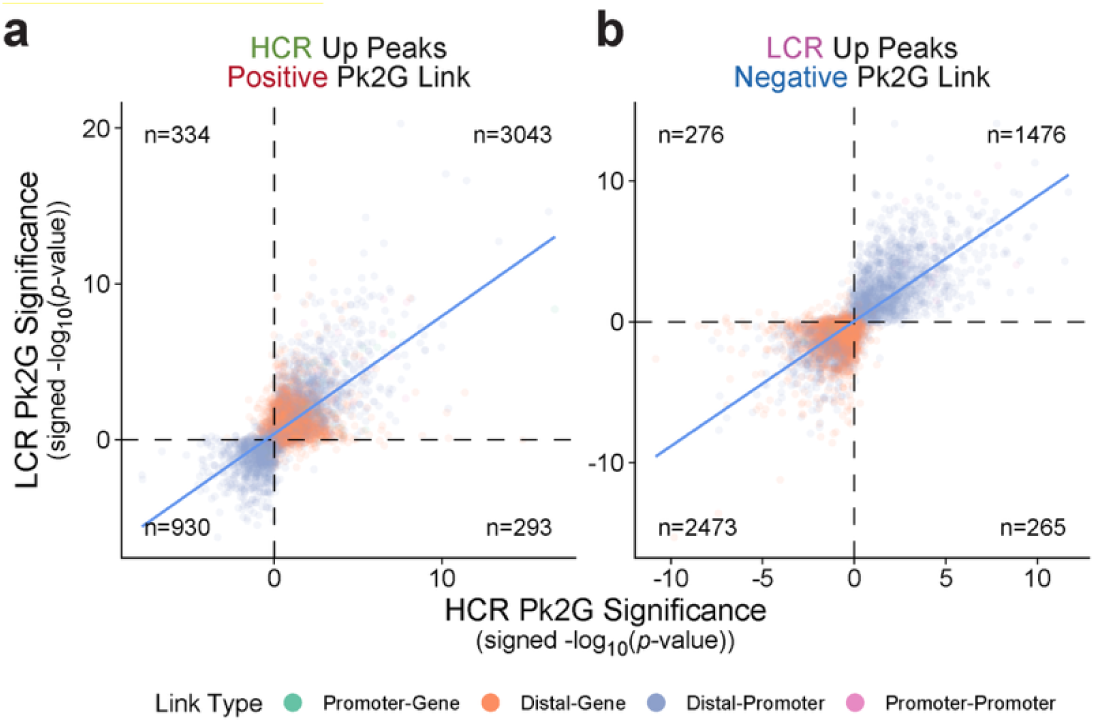
Line-specific Pk2G links, related to Fig. 3. Comparison of line-specific Pk2G links enriched for metabolic pathways in the HCR up positive Pk2G (**a**) and LCR up negative Pk2G (**b**) pathway enrichment analysis. Signed -log_10_(*p*-values) are concordant between the HCR only (x-axis) and LCR only (y-axis) Pk2G analysis. Points are colored by the link type to detail negative and positive distal-promoter raw correlations contributing towards positive and negative Pk2G links, respectively.

**Fig. S11:**
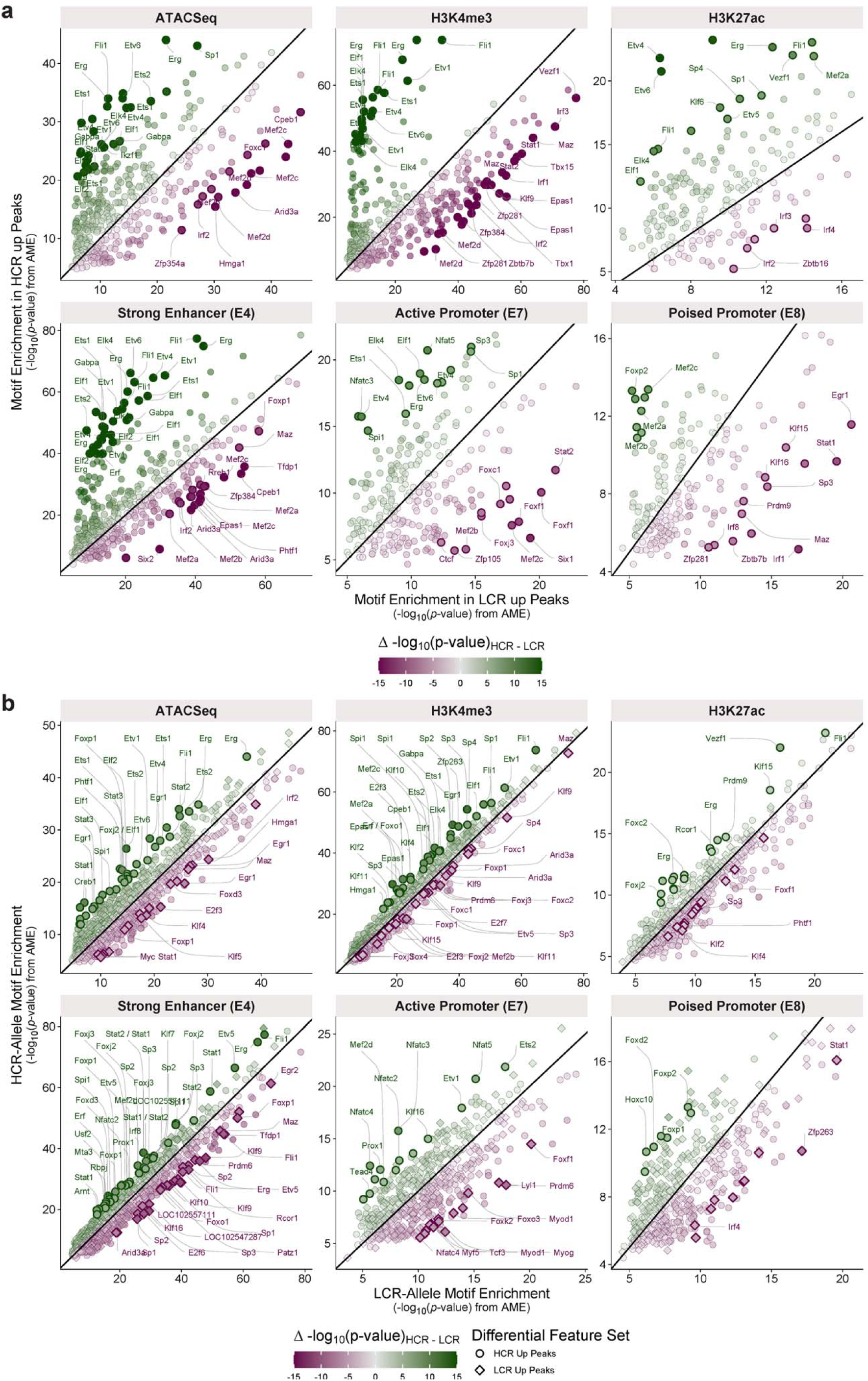
Differential Motif Enrichment, related to Fig. 4. Comparison of motif enrichment from the (**a**) global and (**b**) allele-specific differential motif enrichment analysis across HCR up and LCR up peaks, colored in the same manner as in Fig. 4b,e.

**Fig. S12:**
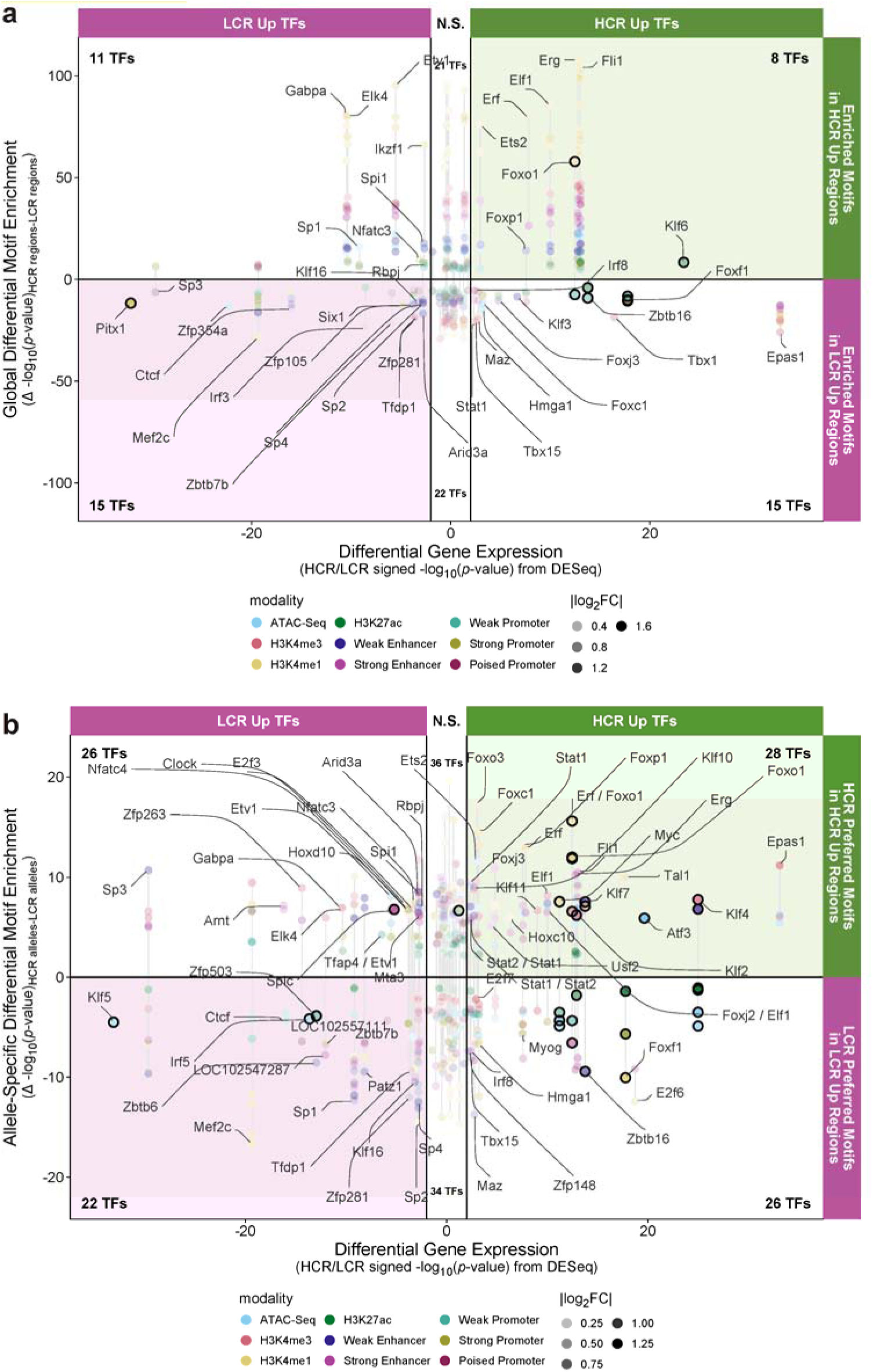
Relationship Between Differential Motifs and Transcription Factor Gene Expression, related to Fig. 4. Comparison of the differential motif enrichment significance (signed Δ -log_10_(AME *p*-value)) across all modalities to the significance of differential TF gene expression (HCR/LCR). The significance of differential gene expression is represented as signed -log_10_(FDR) from DESeq. TFs that are differentially enriched in multiple modalities are connected by a vertical gray line, indicating multiple AME *p*-values with consistent DESeq significance. Black vertical lines at -2 and 2 correspond to a DESeq FDR threshold of < 0.01. The fill intensity represents the magnitude of change (|log_2_FC|). Points with |log_2_FC| > 0.5 are outlined in black. (**a**) Comparison specific to the global motif enrichment analysis. (**b**) Comparison specific to the allele-specific analysis.

**Fig. S13:**
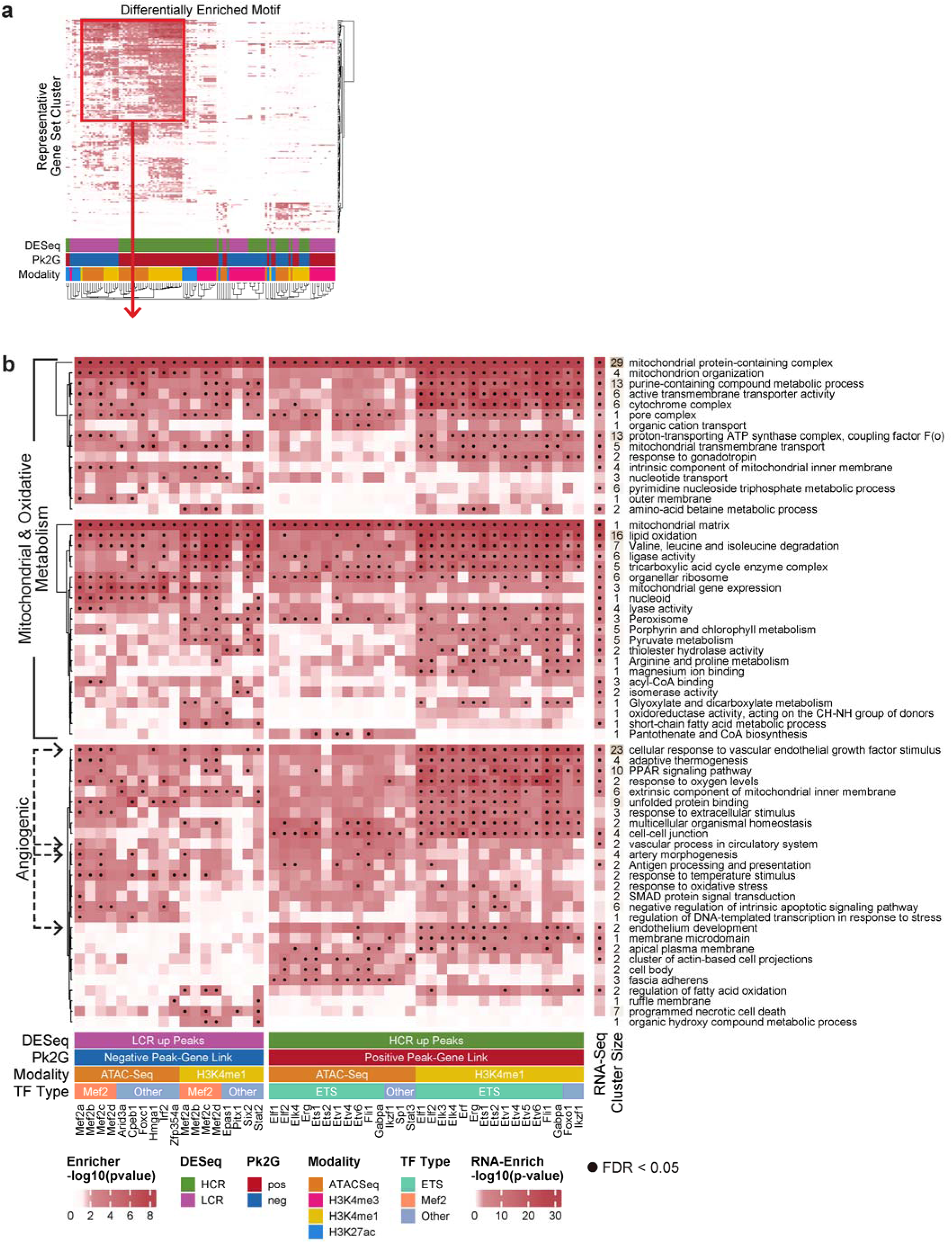
Global HCR/LCR Differential Motif Enrichment Analysis, related to Fig. 4. (**a**) Over-representation analysis of genes linked to differentially enriched motifs outlined in Fig. 4b and Fig. S11a returned 905 significant gene sets (FDR < 0.05) predominantly driven by motifs localized to differential ATAC-Seq and H3K4me1 peaks. The heatmap plots the -log_10_(*p*-values) across analyses stratified by motif, modality, direction of peak change (HCR up or LCR up), and directionality of Pk2G links. Each row is a representative gene set cluster into which significant gene sets (FDR < 0.05) were kappa-reduced. Heatmap presents motifs collapsed into TFs with columns kappa-clustered by gene target similarity and rows hierarchically clustered by -log_10_(*p*-value). Distinct regulatory patterns result, with the most prominent signal driven by motifs enriched in HCR up ATAC-Seq and H3K4me1 peaks, outlined in red. See panel (**b**) for an expanded view of this cluster. (**b**) Subset of the over-representation results from panel (**a**), including the top 10 gene set clusters per column. The number of child gene sets within the representative cluster is annotated on the left-hand side of the heatmap. Results are compared to RNA-Enrich (right column) and the heatmap plots the - log_10_(*p*-value) of the most significant gene set within the cluster with ‘•’ annotating significant clusters (FDR < 0.05). Rows are split into three clusters based on kappa similarity of genes, and then hierarchically by - log_10_(*p*-value) within cluster; columns are manually ordered by column annotation categories. To reduce sparsity in the heatmap, rows are filtered to include terms that are FDR significant in more than 5 ‘motif, modality, HCR/LCR direction, and Pk2G direction’ analyses.

**Fig. S14:**
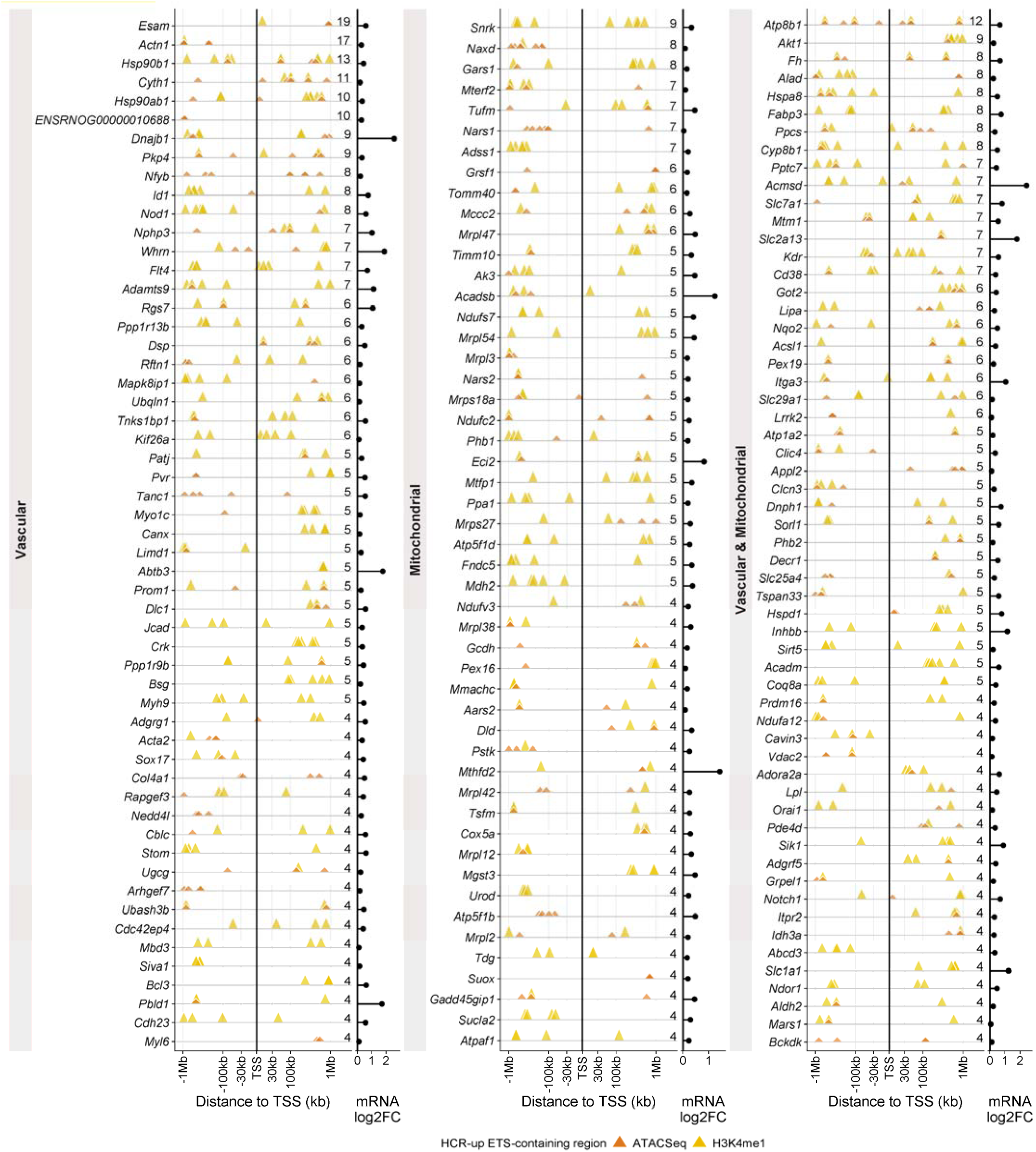
Recurrent ETS-motif regulatory architecture at vascular and mitochondrial genes, related to Fig. 4. Each row is a gene with positive Pk2G links differentially enriched for ETS-family motifs. Triangles represent peaks relative to the gene’s TSS (pseudo-log scale, upstream to the left) and are colored by modality, with the number of linked ETS-containing regions printed on each row. Genes are grouped by the gene set cluster that returned them (Fig. S13). HCR/LCR log_2_FC for each gene is provided on the right.

**Fig. S15:**
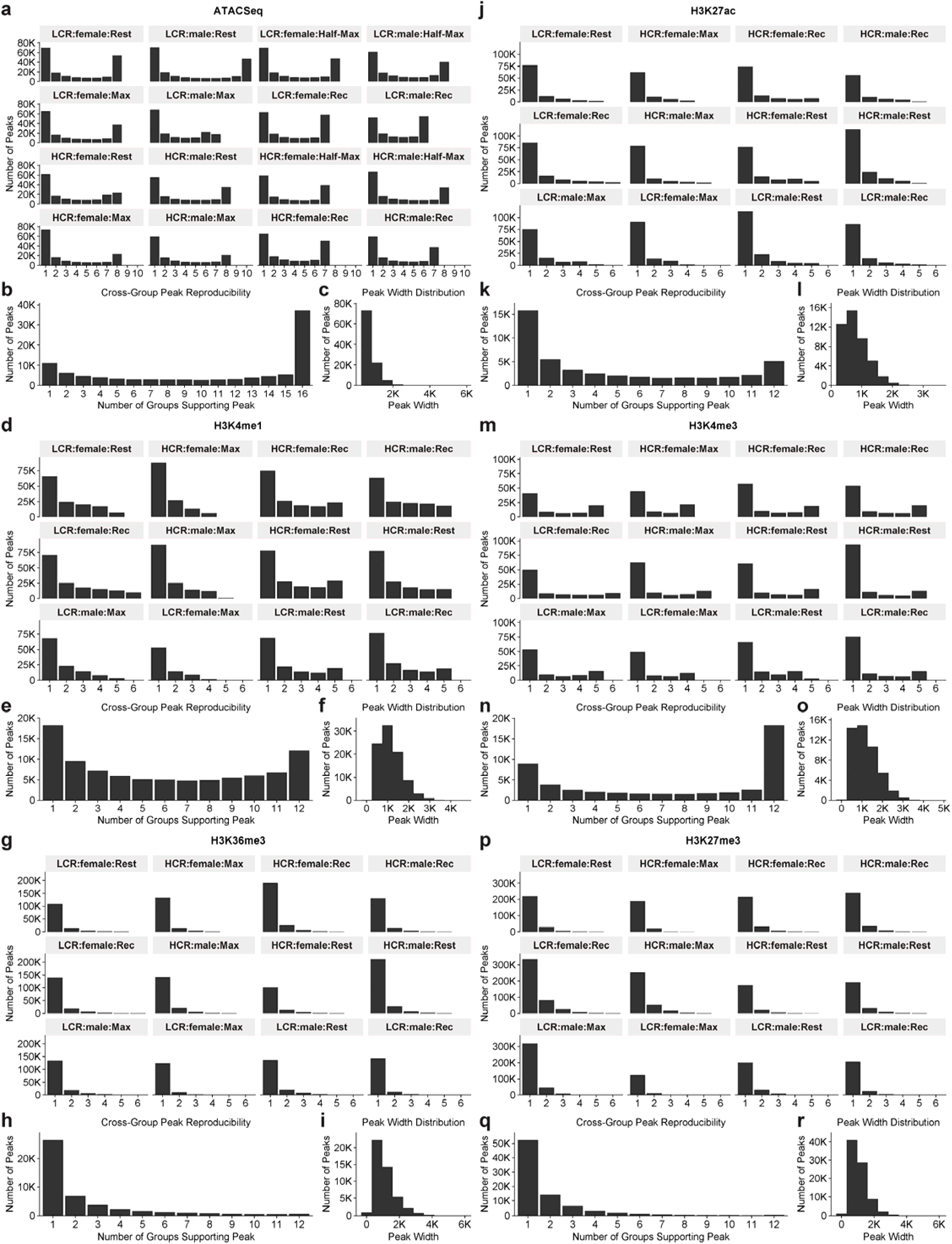
QC Statistics for the Creation of Master Peak Matrices, related to Fig. 2. (**a-r**) Within-group and cross-group reproducibility of peaks called by MACS2, considered when constructing the master peak matrix for each epigenomic sequencing modality.

**Fig. S16:**
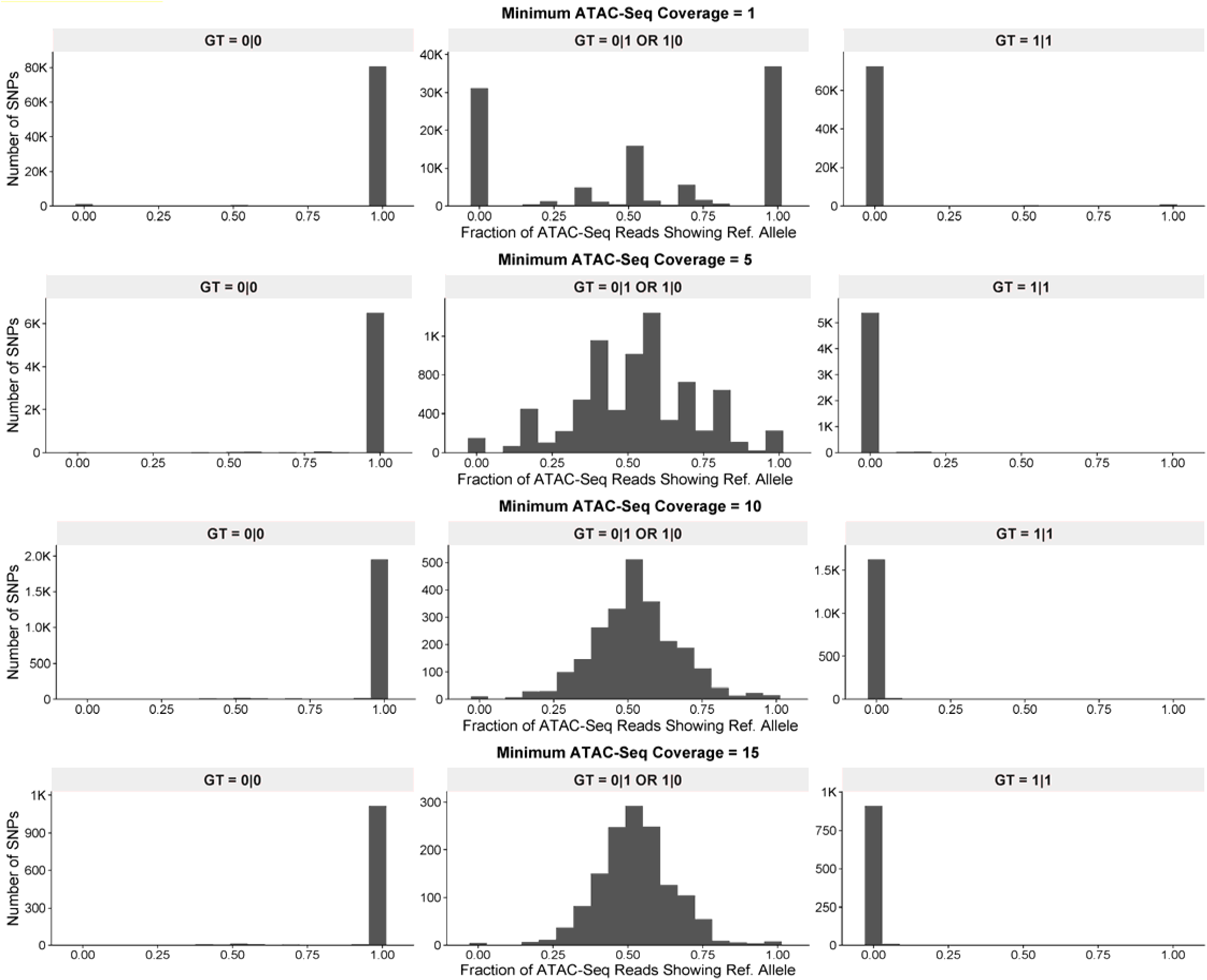
Genotype Imputation QC, related to Fig. 5. Distribution plots showing the proportion of reference alleles present in ATAC-Seq data at variant sites imputed as homozygous reference (0|0), heterozygous (1|0 or 0|1), and homozygous alternate (1|1) at varying ATAC-Seq depth thresholds. Reference alleles are predominantly observed at sites imputed as homozygous reference, and vice versa at sites imputed as homozygous alternate. For sites imputed as heterozygous, the reference allele fraction is centered around 0.5.

## Notes

https://doi.org/10.5281/zenodo.19699175

### Competing Interest Statement

The authors have declared no competing interest.

### Summary of Updates

Title change; Edited abstract; Reorganization of content, including movement of QC details to the Methods and Supplement; Condensed Results; Referred to one consistent result throughout the paper for consistency, rather than introducing new genes to show reproducibility

