## Supplemental Protocol S1 for "Skeletal muscle enhancer programming of intrinsic cardiorespiratory fitness"

**Supplemental Protocol S1: Nuclei Isolation**

Materials
- 2ml glass tissue grinder and pestle ((Kimble chase, 885301-0002, 885300-0002)
- 40 µm cell strainer (Fisher, 22363547)
- Illumina Nextera Tagment DNA enzyme (15027916) (TDE)
- Illumina Tagment DNA buffer (2X) (15027866) (TD buffer)
- NEBNext High-Fidelty 2xPCR Master Mix (NEB M0514S)
- 10,000 x SYBR Green I (Invitrogen Cat# -7563)

| *LB1 buffer* | For 50 ml | For 5ml | For 15 ml | *Final* |
|---|---|---|---|---|
| 1 M HEPES, pH 7.5 | 2.5 ml | 0.25 ml | 0.75 ml | 50mM |
| 5 M NaCl | 1.4 ml | 140 µl | 420 µl | 140 mM |
| 0.5M EDTA, pH 8.0 | 100 µl | 10 µl | 30 µl | 1 mM |
| 50% glycerol | 10 ml | 1 ml | 3 ml | 10% |
| NP-40 10% | 2.5 ml | 0.25 ml | 0.75 ml | 0.5% |
| Triton X-100 10% | 1.25 ml | 125 µl | 375 µl | 0.25% |
| DDW | 32.25 ml | 3.225 ml | 9.675 ml | |
| Immediately before use, add EDTA-free complete mini protease inhibitors (Roche). 1 tablet for each 5 mL buffer. | | | | |

Nuclei extraction
1. Crush frozen tissue (5-10mg) into fine powder while cold (dry ice and LN2).
2. Suspend pulverized tissue in 1 ml ice-cold 1x PBS (1.5 ml tube) by tapping, then centrifuge at 2000g for 3 min. at 4°C.
3. Remove supernatant and resuspend pellet in 1 ml LB1 (1.5 ml tube).
4. Rock tubes at 4°C for 10 min. (Prepare 5 times sample number of 1.5 ml tube if you are doing triplicates, one time the sample number of 2 ml tube, and a 6-well plate for later use)
5. Transfer the sample to a 2 ml glass funnel, loosen stroke (A) 15 times, and then transfer to a 1.5 ml tube.
6. Centrifuge at 2000g for 5 min at 4°C.
7. Aspirate supernatant and resuspend in 1 ml 1x cold PBS.
8. Filter with a 40-µm cell strainer (in a 6-well plate), and wash the plate with 1 ml of 1x PBS.
9. Count nuclei with a cell counter (ideally want 10^5 nuclei; tryptan blue stains nuclei; typically 4-9 µm)
10. Transfer 50,000 nuclei (mix before transfer) to a 1.5 ml tube, and replicate or triplicate each sample here. If the volume is below 50 µL, add PBS to a 1.5 ml tube to make the volume 50 µl. Centrifuge at 500 x g for 10 min at 4 °C (be careful of the tube direction). Aspirate supernatant with pipette (should be very careful).
